# Therapeutic KRAS^G12C^ inhibition drives effective interferon-mediated anti-tumour immunity in immunogenic lung cancers

**DOI:** 10.1101/2021.10.18.464819

**Authors:** Edurne Mugarza, Febe van Maldegem, Jesse Boumelha, Christopher Moore, Sareena Rana, Miriam Llorian Sopena, Philip East, Rachel Ambler, Panayiotis Anastasiou, Pablo Romero Clavijo, Karishma Valand, Megan Cole, Miriam Molina-Arcas, Julian Downward

**Affiliations:** Oncogene Biology Laboratory, Francis Crick Institute, 1 Midland Road, London NW1 1AT, UK; Bioinformatics and Biostatistics Science Technology Platform, Francis Crick Institute, 1 Midland Road, London NW1 1AT, UK; Lung Cancer Group, Division of Molecular Pathology, Institute of Cancer Research, 237 Fulham Road, London SW3 6JB, UK

## Abstract

Recently developed KRAS^G12C^ inhibitory drugs are beneficial to lung cancer patients harbouring KRAS^G12C^ mutations, but drug resistance frequently develops. Due to the immunosuppressive nature of the signaling network controlled by oncogenic KRAS, these drugs can indirectly affect anti-tumour immunity, providing a rationale for their combination with immune checkpoint blockade. In this study, we have characterised how KRAS^G12C^ inhibition reverses immune suppression driven by oncogenic KRAS in a number of pre-clinical lung cancer models with varying levels of immunogenicity. Mechanistically, KRAS^G12C^ inhibition upregulates interferon signaling via Myc inhibition, leading to reduced tumour infiltration by immunosuppressive cells, enhanced infiltration and activation of cytotoxic T cells, and increased antigen presentation. However, the combination of KRAS^G12C^ inhibitors with immune checkpoint blockade only provides synergistic benefit in the most immunogenic tumour model. KRAS^G12C^ inhibition fails to sensitize cold tumours to immunotherapy, with implications for the design of clinical trials combining KRAS^G12C^ inhibitors with anti-PD1 drugs.

**One sentence summary:** KRAS inhibition mobilizes anti-tumour immunity in immunogenic lung cancer models through derepressing interferon signaling via repression of Myc.

## INTRODUCTION

Lung cancer is the number one cause of cancer deaths worldwide, leading to some 1.8 million deaths annually, and therefore represents a disease of very high unmet need (*1*). Non-small cell lung cancer (NSCLC) comprises 84% of all lung cancers and has a 5-yr survival rate of only 25% (*2*). Fortunately, with the introduction of immune checkpoint blockade (ICB), such as anti-PD1 therapy, aiming to boost anti-tumour T cell immunity, the paradigm for treatment has shifted, enabling long-lasting responses in a subset of patients (*3*). However, only a minority of patients respond and, of those that do, many develop resistance to treatment over time, hence great efforts are currently aimed at trialing therapeutic combinations with ICB (*4*). Targeting oncogenic drivers has been another approach to control tumour growth, as recurrent genetic alterations are detected in more than half of lung adenocarcinoma patients (*5*).

Targeted inhibition of receptor tyrosine kinases such as EGFR has extended progression-free survival beyond conventional cytotoxic therapies. But, until recently, inhibiting KRAS, the most frequent target of oncogenic mutations found in about 15% of all cancer patients and 33% of those with lung adenocarcinoma (*6*), has been notoriously difficult. In 2013, Ostrem et al. reported the development of a covalent inhibitor that was able to lock KRAS into its inactive GDP-bound state by binding to the cysteine resulting from the G12C mutation, present in 40% of KRAS-mutant NSCLC patients (*7*). A mutation-specific inhibitor would be able to circumvent the high toxicity that has limited the widespread use of compounds targeting signaling downstream of KRAS, such as MEK inhibitors. The discovery of a KRAS^G12C^-specific compound led to rapid development of clinical inhibitors and in 2021 Amgen was the first to obtain FDA approval for clinical use of AMG510 (Sotorasib) in locally advanced or metastatic KRAS^G12C^-mutant NSCLC (*8–10*). As expected, toxicity from these drugs is low as signaling is only inhibited in cancer cells harboring the G12C mutation, and clinical response rates are high, but unfortunately resistance occurs frequently within a few months of treatment. Several mechanisms of resistance have already been described, including novel mutations in KRAS or bypassing the mutation via redundant signaling pathways (*11–13*). Hence, combination therapies will be needed to make a greater impact on patient survival (*14, 15*).

Exploring the combination of targeted inhibition of KRAS with anti-PD1 therapy seems an obvious approach and indeed the first clinical trials are already well underway. There are certainly rational arguments to make for this combination, with KRAS mutant lung cancer generally being associated with a smoking-history and therefore high tumour mutation burden, one of the positive predictors for response to ICB (*16, 17*). Moreover, KRAS-mutant lung cancer is strongly associated with an immune evasive phenotype and KRAS signaling is thought to play a role in orchestrating such an immune suppressive environment, for example by driving the expression of cytokines and chemokines as was shown for IL-10, TGF-beta and GM-CSF in KRAS mutant pancreatic cancer (*18*). Inhibition of KRAS could provide temporary relief from such immune suppression and a window of opportunity for T cell activation. Indeed, initial reports of KRAS^G12C^ inhibition showed that durable responses in mice were dependent on T cells, and a combination of KRAS^G12C^ inhibition and anti-PD1 led to improved survival in a subcutaneous tumour model of the genetically engineered G12C KRAS mutant CT26 colon cancer cell line (*8, 19*). While the KRAS^G12C^ mutation is only found in 3-4% of colon carcinoma, it is more prevalent in NSCLC (∼14%) and clinical efficacy of the KRAS inhibitors also seems to be higher in lung cancer. Therefore, it will be most relevant to understand the mechanisms underlying potential therapeutic cooperation between KRAS inhibition and immune responses in the setting of lung cancer (*9*). Tissue site, existing immune evasive tumour microenvironment (TME) and intrinsic immunogenicity in the form of neoantigen presentation are all likely to be important factors in determining the outcome of combination treatments with ICB (*4*). Fedele, et al. showed that a combination of SHP2 and KRAS^G12C^ inhibition led to good tumour control and increased T cell infiltration in an orthotopic model of lung cancer (*20*). Using the strongly immune evasive 3LL *Δ*NRAS lung cancer cell line (*14*), we recently developed an imaging mass cytometry analysis pipeline which showed that KRAS^G12C^ inhibition was able to induce remodeling of the lung TME (*21*). Here, we use this 3LL *Δ*NRAS alongside other pre-clinical lung cancer models varying in degree of immunogenicity to perform an in-depth investigation of the impact of KRAS^G12C^ inhibition on the TME and anti-tumour immunity and explore the mechanisms that underly the changes observed. We describe several mechanisms by which tumour-specific KRAS inhibition has direct and indirect effects on the TME, such as reduced expression of chemokines attracting immune suppressive myeloid cells, enhanced uptake of tumour cells by antigen presenting cells, and enhanced intrinsic and extrinsic interferon (IFN) responses. Furthermore, we show that successful combination of KRAS^G12C^ inhibition with ICB is not universal, but rather varies between the models and correlates with immunogenicity, which will have important implications for the selection of patients that may benefit from such combination therapy. In particular, tumours that are refractory to ICB, either due to intrinsic or acquired resistance, may be unlikely to be resensitised by combination with KRAS^G12C^ inhibition alone.

## RESULTS

### Oncogenic KRAS regulates expression of cytokine and immune regulatory genes in human and murine cell lines

Previous reports have described that KRAS signaling can mediate the expression of cytokines such as IL-8 and GM-CSF in pancreatic cancer models (*22*). We therefore decided to assess the role of oncogenic KRAS signaling in the regulation of the expression of immunomodulatory factors in the lung. For this purpose, we made use of a cell line model of immortalised human lung pneumocytes expressing a tamoxifen inducible oncogenic KRAS protein (KRAS^G12V-ER^) (*14*) (Fig S1A). Activation of oncogenic KRAS signaling induced the secretion of a number of cytokines and chemokines which could affect the recruitment and polarisation of different immune cells (Fig 1A). To further expand the scope of our investigation we performed whole transcriptome analysis (RNA-Seq) on this model. Results validated the transcriptional induction of myeloid cell modulatory factors IL-8, CXCL1, CCL2, GM-CSF, and several other KRAS regulated cytokines (Fig 1B). Additionally, gene set pathway analysis revealed a KRAS-dependent negative regulation of type I and type II interferon (IFN) responses in lung pneumocytes (Fig 1C and S1B), previously shown to be crucial for anti-tumour immunity and sensitivity to immunotherapy (*23, 24*) which could reflect a mechanism triggered by oncogenic KRAS to promote immune evasion.

**Fig. 1.**
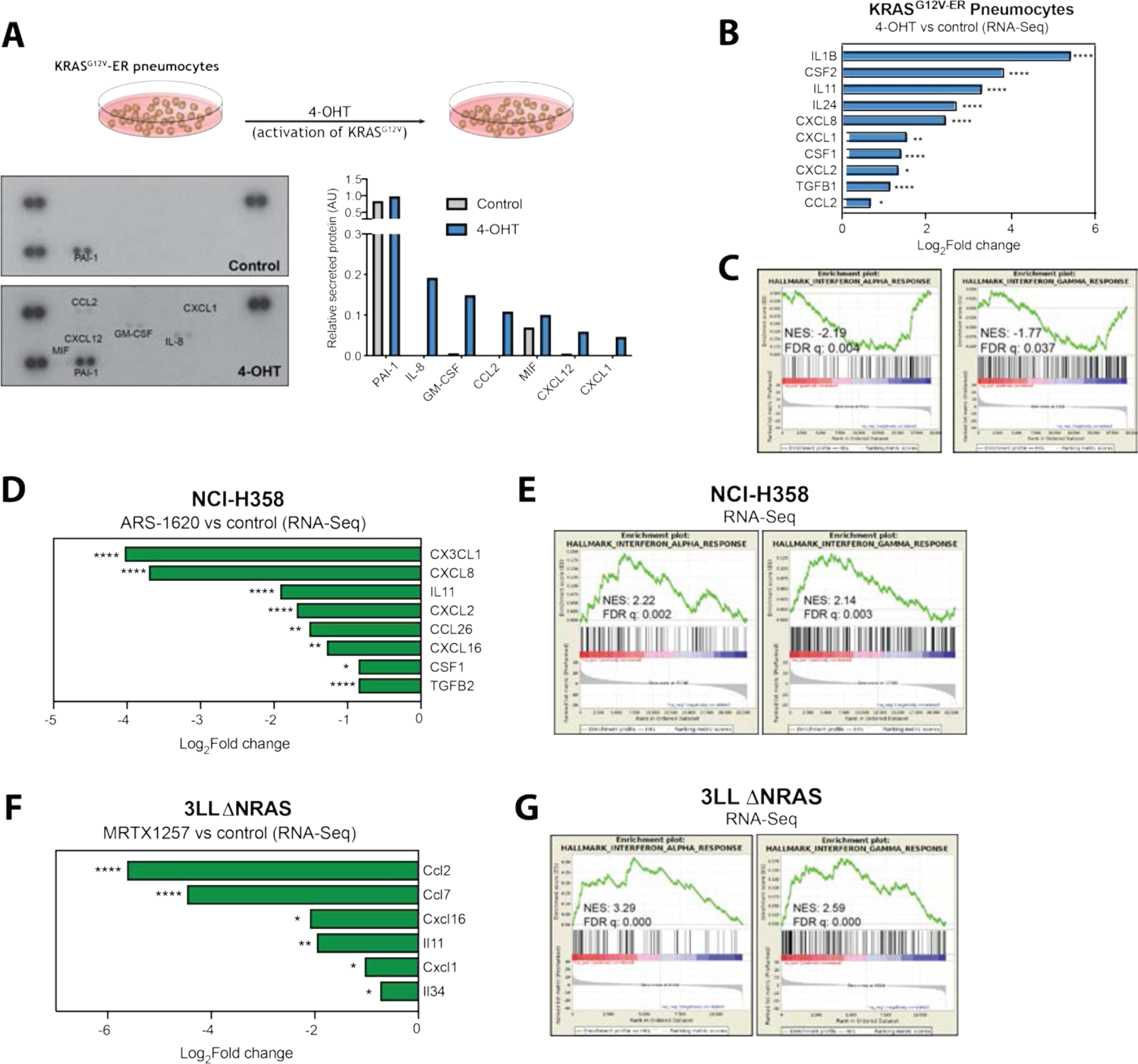
Oncogenic KRAS regulates immune gene expression in cell lines. (A) Cytokine array of cell culture supernatant from KRAS^G12V-ER^ pneumocytes treated with 500nM 4-OHT or ethanol control for 24h. Graph shows secreted protein relative to control spots on the array for each condition. (B) Log_2_Fold change of selected cytokine genes from RNA-Seq data in 4-OHT (500nM, 24h) treated KRAS^G12V-ER^ pneumocytes. (C) MSigDB Hallmarks gene set enrichment analysis (GSEA) plots of IFN*α* and IFN*γ* pathways in 4-OHT treated versus control samples. (D) Log_2_Fold change of selected cytokine genes from RNA-Seq data in ARS-1620 (2μM, 24h) treated NCI-H358 cells versus DMSO control. (E) MSigDB Hallmarks GSEA plots of IFN*α* and IFN*γ* pathways in ARS-1620 treated versus control samples. (F) Same analysis as (D) of RNA-Seq from 3LL *Δ*NRAS cells treated with 100nM MRTX1257 (24h, n=3). (G) Same analysis as (E) of RNA-Seq from 3LL *Δ*NRAS cells. All statistics represent FDR adjusted p values (q<0.05).

We then assessed whether treatment with a therapeutic KRAS^G12C^ inhibitor could reverse these mechanisms. In two KRAS^G12C^-mutant human lung cancer cell lines, abrogation of oncogenic KRAS signaling by a KRAS^G12C^ inhibitor (Fig S1C) led to the downregulation of cytokines and chemokines, particularly those involved in the recruitment and differentiation of myeloid cell populations, known to exert tumour-promoting effects in the tumour microenvironment (Fig 1D and S1D top). Interestingly, only the neutrophil chemoattractants CXCL2 and CXCL8 were consistently KRAS-regulated in both models, while most factors were cell line-specific, suggesting that different cell lines exhibit different cytokine expression patterns. RNA-Seq analysis of these cell lines also revealed that KRAS^G12C^ inhibition upregulates IFN*α* and IFN*γ* response gene expression, a mechanism that was consistent across both cell lines (Fig 1E and S1D bottom).

We decided to extend our findings to a murine cell line in order to use immunocompetent mouse models to examine the effects of oncogenic KRAS on anti-tumour immunity in vivo. We made use of a murine transplantable KRAS^G12C^-mutant lung cancer cell line derived from Lewis Lung Carcinoma, 3LL *Δ*NRAS (described in (*14*)), which is sensitive to KRAS^G12C^ inhibition (Fig S1E). Using this model, we validated the effect of KRAS^G12C^ inhibitors on the transcriptomic downregulation of secreted immunomodulatory factors, by both RNA-Seq (Fig 1F) and qPCR (Fig S1F) analysis. Likewise, we validated the upregulation of type I and II IFN gene sets observed in previous models (Fig 1G and S1G).

Together, these data suggest that oncogenic KRAS signaling regulates the expression of factors that could affect the tumour microenvironment and anti-tumour immunity and highlight the role of KRAS^G12C^ inhibitors in reversing these potentially immune evasive mechanisms.

### KRAS signaling downregulates IFN pathway gene expression via MYC

Next, we decided to further investigate the mechanistic link between KRAS signaling and IFN responses given their important role in anti-tumour immunity. We began by validating our RNA-Seq finding that genes coding for components of the IFN response were upregulated by KRAS^G12C^ inhibition in 3LL *Δ*NRAS cells (Fig 2A). This upregulation occurred in a MEK-dependent and cell viability-independent manner, beginning at approximately 6 hours after treatment and peaking at 24 hours after treatment (Fig S2A-C). To extend our findings we used two additional mouse cell lines modified to harbor KRAS^G12C^ mutations, the KRAS mutant, p53 deleted lung cancer cell line KPB6^G12C^ (*25*) and the KRAS mutant colon cancer cell line CT26^G12C^ (*19*). In these models, treatment with KRAS^G12C^ inhibitor MRTX1257 (MRTX) consistently led to the upregulation of canonical IFN signaling pathway genes (Fig 2B). Interestingly, MEK inhibition in non-G12C mutant isogenic KPB6 cell lines also increased IFN signaling gene expression (Fig S2D), indicating that the mechanism is conserved across oncogenic KRAS mutations.

**Fig. 2.**
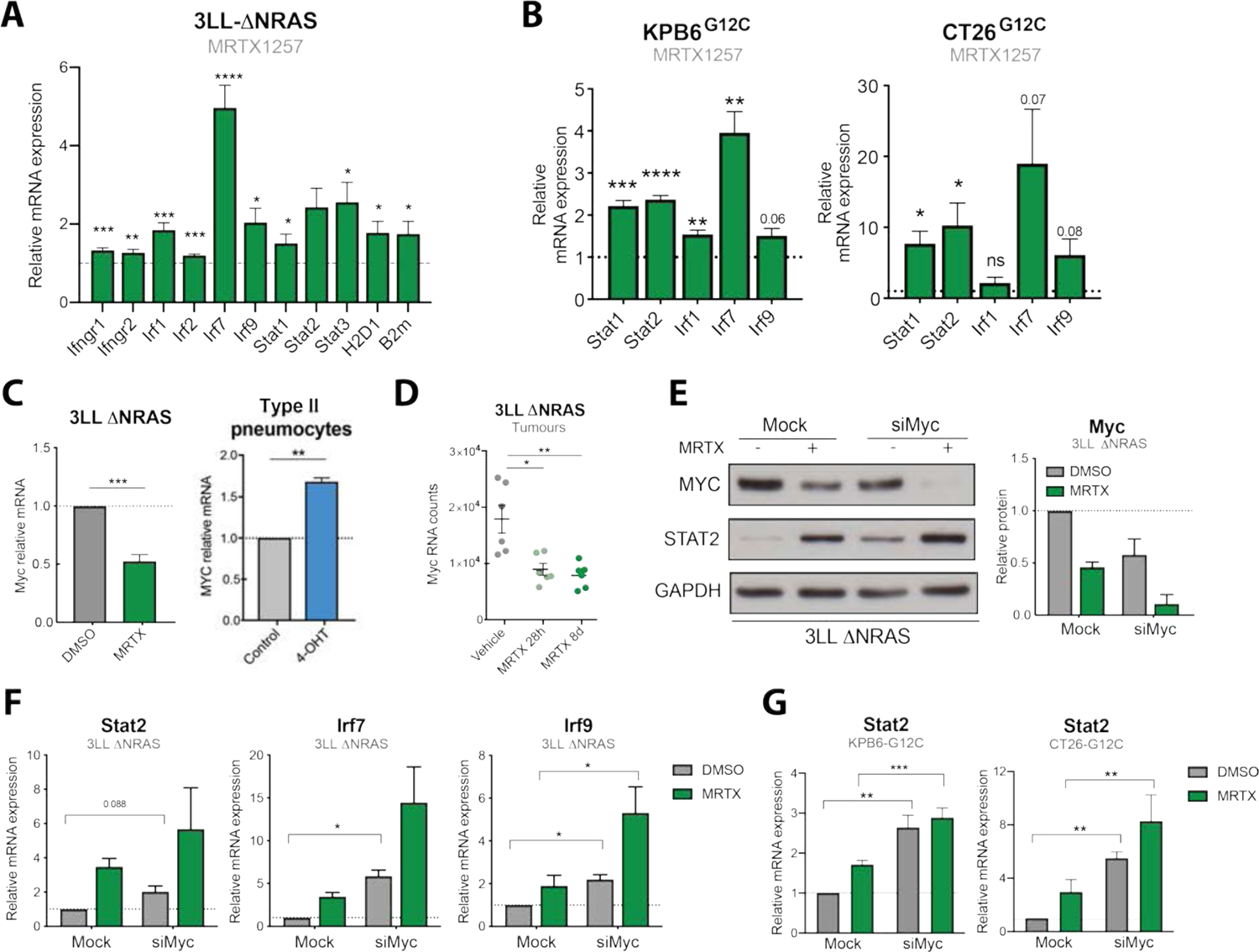
KRAS signaling downregulates IFN pathway gene expression via MYC. (A) qPCR analysis of IFN-induced genes in MRTX1257-treated (100nM, 24h) 3LL *Δ*NRAS cells (2^-ΔΔCT^, normalised to control sample for all genes, n=6, unpaired t test, mean+SEM). (B) Same as (A) using KPB6^G12C^ (n=4) and CT26^G12C^ (n=3) cell lines. (C) qPCR showing KRAS-dependent regulation of Myc in 3LL *Δ*NRAS cells (n=3) after treatment with MRTX1257 and KRAS^G12V-ER^ pneumocytes (n=4) after treatment with 4-OHT (unpaired t test, mean+SEM). (D) RNA-Seq mRNA counts of 3LL *Δ*NRAS lung tumours treated with vehicle or 50mg/kg MRTX1257 for 28h or 8 days (each dot represents a tumour, n=6 per group, FDR p adjusted value). (E) Western blot showing MYC knockdown and STAT2 upregulation of 3LL *Δ*NRAS cells treated with 100nM MRTX1257 (24h), Myc siRNA (48h), or both. Quantification for two independent experiments is shown on the right (mean+SEM). (F) qPCR analysis of IFN-induced genes in 3LL *Δ*NRAS cells treated with 100nM MRTX1257, Myc siRNA or both (2^-ΔΔCT^, normalised to control sample for all genes, n=3, paired t tests siMyc versus Mock, mean+SEM). (G) Same analysis as (F) in KPB6^G12C^ (n=4) and CT26^G12C^ (n=3) cells.

The conserved regulation of IFN signaling pathway genes after KRAS inhibition could suggest a direct crosstalk between oncogenic KRAS and IFN signaling pathways. IFNs bind their receptors on the membrane of target cells and drive transcriptional changes via activation of JAK-STAT signaling modules. To investigate whether the increase in gene expression in response to KRAS inhibition was a result of augmented IFN signaling, we examined the effect of blocking or depleting individual IFN pathway components. However, antibody-mediated blocking of the IFN*α*receptor (Fig S2E), pharmacological inhibition of JAK1/2 signal transduction with ruxolitinib (Fig S2F) and gene knockdown of Stat1 or Stat2 (Fig S2G) did not affect the MRTX-driven upregulation of IFN genes (Fig S2H), suggesting that KRAS-driven inhibition of the IFN pathway occurs independently of the interferon receptors and JAK-STAT proteins.

A known negative transcriptional regulator of IFN genes is the MYC oncoprotein, which is also a RAS target (*26*). As expected, MYC mRNA levels were downregulated by KRAS^G12C^ inhibition in vitro and in vivo and upregulated after KRAS^G12V^ activation (Fig 2C and D), confirming the KRAS-driven regulation of MYC in our models. We assessed the role of MYC in the regulation of IFN signaling pathway genes by the KRAS^G12C^ inhibitor. In the 3LL *Δ*NRAS cell line, despite incomplete knockdown (Fig 2E), MYC depletion was able to significantly increase the expression of these genes (Fig 2F). Because MRTX treatment led to a further downregulation of MYC, increased gene expression was observed when MRTX and siMyc were combined. Importantly, upregulation of IFN signaling pathway genes in response to MYC depletion was a common response observed across the three murine KRAS^G12C^ cell lines (Fig S2I and S2J). Furthermore, in CT26^G12C^ and KPB6^G12C^ cells, where near-complete knockdown of MYC was achieved (Fig S2I), no significant additional effects on gene or protein expression were observed by combined siMyc and MRTX treatment (Fig 2G), suggesting that MRTX-driven regulation of these genes is primarily through MYC. Together, these data suggests that KRAS^G12C^ inhibition, through downregulation of MYC, leads to increased expression of genes associated with the IFN response.

### KRAS^G12C^ inhibition enhances tumour cell intrinsic IFN responses

We next investigated whether KRAS^G12C^ inhibitor-driven changes in gene expression affected the capacity of tumour cells to respond to IFN*γ*. We found that, indeed, IFN*γ*-driven transcriptional effects were enhanced by MRTX treatment (Fig 3A). KRAS^G12C^ inhibition augmented the IFN*γ*-driven expression of immunomodulatory IFN-stimulated genes (ISG) such as T cell chemoattractants *Cxcl9/10/11* and antigen presentation genes including *H2-d/k1*, *Ciita* and *B2m* (Fig 3A and S3A). Consistent with the KRAS-dependent regulation of type I and II IFN responses observed in our RNA-Seq analysis, MRTX treatment was likewise able to enhance IFN*α* and IFN*β*-driven gene expression (Fig S3B). These transcriptional changes also led to increased protein expression of ISG, as evidenced by an increased proportion of IFN*γ*-induced CXCL9-secreting tumour cells after treatment with MRTX (Fig 3B). We validated that MRTX treatment enhanced IFN*γ*-driven gene (Fig 3C) and protein (Fig S3C and S3D) expression in two additional murine cell lines. A similar improvement of responses to IFN*γ* could be achieved by MYC knockdown (Fig 3D), confirming the role of MYC in the regulation of IFN responses.

**Fig. 3.**
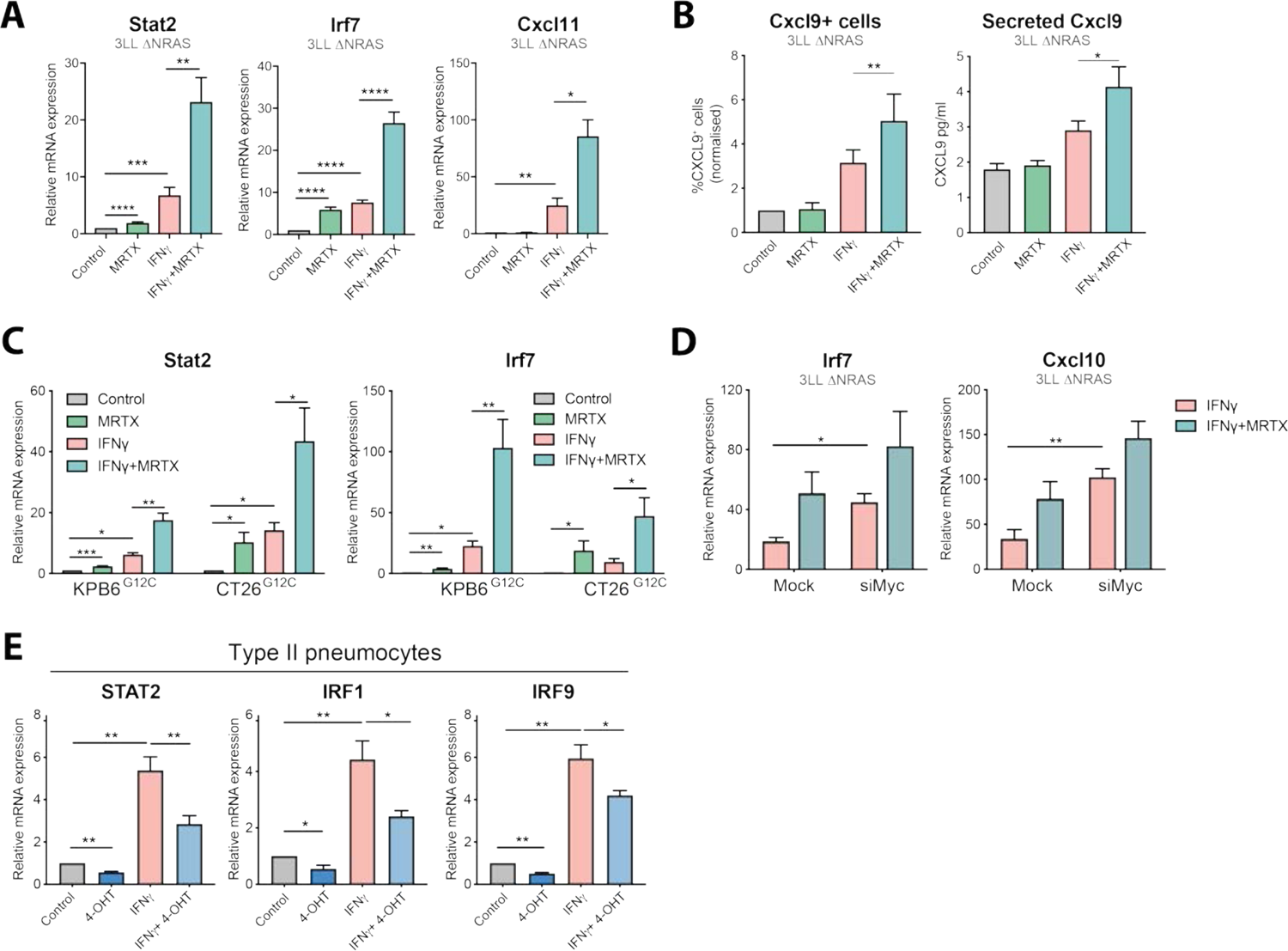
KRAS^G12C^ inhibition enhances tumour cell intrinsic IFN responses. (A) qPCR analysis of IFN-induced genes in MRTX1257 (100nM, 24h) and/or recombinant IFN*γ* (100ng/ml) treated 3LL *Δ*NRAS cells (2^-ΔΔCT^, normalised to control sample for all genes, n=6, paired t test, mean+SEM). (B) Protein validation of IFN response regulation by KRAS. Left: Percentage of CXCL9-positive cells as measured by flow cytometry on 3LL *Δ*NRAS cells after treatment with MRTX1257 and/or IFN*γ*. Right: concentration of CXCL9 secreted to the medium of 3LL *Δ*NRAS cells after treatment with MRTX1257 and/or IFN*γ*, measured by ELISA (normalised to control sample, n=3, paired t test, mean+SEM for both). (C) Same as (A) using KPB6^G12C^ (n=4) and CT26^G12C^ (n=3) cell lines. (D) qPCR analysis of IFN-induced genes in 3LL *Δ*NRAS cells treated with IFN*γ* only or IFN*γ* and MRTX1257 in presence of 48h of Mock or Myc siRNA (2^-ΔΔCT^, normalised to IFN*γ* only-treated sample for all genes, n=3, paired t test, mean+SEM). (E) qPCR analysis of IFN pathway genes in human KRAS^G12V-ER^ pneumocytes after treatment with 4-OHT and/or recombinant IFN*γ* for 24h (normalised to control sample n=4, paired t test, mean+SEM).

To exclude the possibility that reduced cell fitness contributed to the effects of KRAS^G12C^ inhibition on the response to IFN*γ*, we validated our findings in the KRAS^G12V-ER^ human pneumocyte cell line. In this model, we observed that 4-OHT-induced KRAS^G12V^ activation led to the downregulation of IFN signaling pathway genes and was able to decrease IFN*γ*-driven transcriptional effects (Fig 3E), suggesting a mechanistic link between the KRAS and IFN pathways which is not influenced by cell viability.

In summary, we have shown that oncogenic KRAS signaling can suppress responses to IFN, and that this can be alleviated with pharmacological KRAS^G12C^ inhibition. KRAS inhibition in consequence leads to an increased sensitivity of tumour cells to type I and II IFNs, which translates to higher expression of IFN-induced genes such as T cell chemoattractants and antigen presentation genes that could positively affect anti-tumour immunity in vivo.

### KRAS^G12C^ inhibition in vivo remodels the highly immunosuppressive TME of 3LL ΔNRAS lung tumours

The results presented above demonstrate the ability of KRAS^G12C^ inhibitors to reverse KRAS-driven immune evasion mechanisms, such as enhancing tumour cell-intrinsic IFN responses and modulating the expression of secreted immunomodulatory factors. Next, we aimed to assess how inhibition of oncogenic KRAS in vivo can affect the composition of the tumour microenvironment (TME) in lung tumours.

The 3LL *Δ*NRAS cell line can form orthotopic tumours in the lungs of C57BL/6 mice when delivered intravenously. Immunophenotypic characterisation of these tumours revealed a predominant infiltration of myeloid cells, known to exert immunosuppressive actions, while anti-tumorigenic cells like lymphocytes and NK were largely absent from the TME (Fig 4A). Using Imaging Mass Cytometry (IMC) we recently showed that there was an inclusion of macrophages and neutrophils in the core of 3LL *Δ*NRAS lung tumours, while effector cells remained at the tumour periphery (*21*). Consistent with this apparent immunosuppressive TME, growth of these tumours was not affected by a lack of B and T cells in Rag1^-/-^ mice (Fig S4A). Whole exome sequencing of two 3LL *Δ*NRAS single cell clones derived from the CRISPR-Cas9 deletion of NRAS (*14*) revealed that these cell lines harbor thousands of clonal somatic nonsynonymous single nucleotide variations (SNVs) compared to the reference C57BL/6J genome (Fig 4B). Whole exome sequencing and RNA expression data were combined to perform in silico neoantigen prediction. Results showed that this cell line harbors numerous predicted neoepitopes with high or medium affinity for MHC binding (Fig S4B) but flow cytometric analysis revealed that it has lost the expression of one of the MHC alleles, H2-Kb, while retaining the other, H2-Db (Fig S4C), possibly reflecting a mechanism to escape immunological rejection. It is noticeable that neoantigens predicted to bind to the absent H2-Kb are several-fold more highly represented than neoantigens precited to bind to the expressed H2-Db. Therefore, we hypothesize that this cell line contains sufficient neoantigens to elicit an anti-tumour immune response, yet it is highly immune evasive and avoids rejection in immunocompetent hosts, likely through a combination of mechanisms including reduced neoantigen presentation and production of an immunosuppressive tumour microenvironment.

**Fig. 4.**
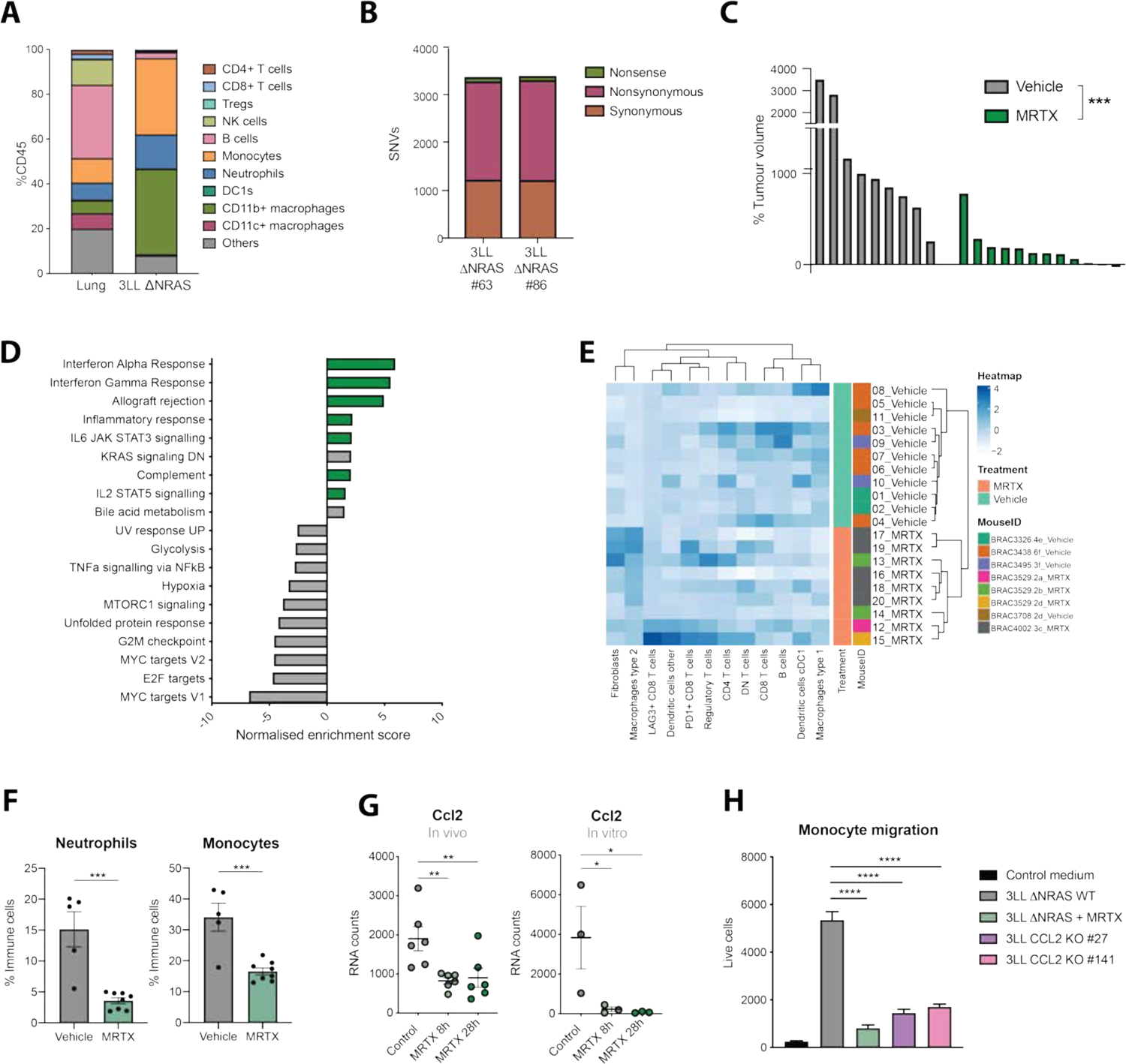
KRAS^G12C^ inhibition remodels the immunosuppressive TME of 3LL ΔNRAS lung tumours. (A) Immunophenotyping of dissected lung tumours obtained by intravenous administration of 3LL *Δ*NRAS cells (n=5 mice) versus healthy lung tissue (n=6 mice) obtained by flow cytometry. (B) Whole exome sequencing SNV analysis of two NRAS CRISPR-edited 3LL clones. (C) Post-treatment tumour volume change as measured by μCT scanning of 3LL *Δ*NRAS lung tumours after one week of treatment with vehicle control or 50mg/kg MRTX1257 (Each bar represents one tumour, Mann-Whitney test). (D) Summary of significantly (FDR q<0.05) up- and down-regulated pathways in MRTX-versus vehicle-treated lung tumours (MSigDB Hallmarks). (E) Hierarchical clustering of relative frequencies of tumour infiltrating cell types in MRTX- and vehicle-treated tumours obtained by IMC. (F) Percentage of neutrophils (gated as CD45+ CD11b+ Ly6C+ Ly6G+) and monocytes (gated as CD45+ CD11b+ Ly6C^hi^ Ly6G-) in vehicle (n=5) and MRTX-treated (n=8) lung tumours measured by flow cytometry (each dot represents a mouse, unpaired t test). (G) mRNA counts for *Ccl2* gene in MRTX treated 3LL *Δ*NRAS tumours (n=6 per group, left) and cells (n=3, right) obtained by RNA-Seq (FDR adjusted p value). (H) Live cell count (by flow cytometry) of bone marrow-derived monocytes that have migrated through a transwell in presence of conditioned medium from 3LL *Δ*NRAS cells, MRTX-treated cells or two clones from *Ccl2* CRISPR knockout (n=3 independent experiments, one-way ANOVA).

Treatment of 3LL *Δ*NRAS lung tumour-bearing mice with the KRAS^G12C^ inhibitor MRTX1257 for one week resulted in marked tumour growth inhibition (Fig 4C), although relatively few tumours actually decreased in size, highlighting the extreme aggressiveness of this tumour model. At this time point, we harvested tumours to perform RNA-Seq analysis, flow cytometric analysis of immune cell infiltration and IMC to examine the effects of KRAS^G12C^ inhibition on the TME of this highly immunosuppressive model. As anticipated from the in vitro data, gene set enrichment analysis of 3LL *Δ*NRAS lung tumours treated with MRTX revealed an upregulation of several immune-related pathways, including interferon *α* and *γ*responses, IL2 and IL6 signaling, allograft rejection, complement and inflammatory responses (Fig 4D). We were likewise able to confirm the KRAS-dependent regulation of IFN signaling pathway gene sets in vivo (Fig S4D). This striking remodeling of the TME was confirmed by IMC analysis, where hierarchical clustering based on immune cell infiltration patterns was able to discern vehicle and MRTX-treated samples (Fig 4E).

KRAS^G12C^ inhibition was able to significantly reduce the high infiltration of myeloid cells like monocytes and neutrophils observed in this lung tumour model, as measured by flow cytometry (Fig 4F). We then wondered whether the downregulation of tumour cell-intrinsic cytokine expression observed in vitro (Fig 1F) could play a role in the regulation of myeloid cell infiltration. The strongest KRAS-regulated cytokine in the 3LL *Δ*NRAS cells in vitro was *Ccl2*, a canonical chemoattractant for monocytes. The KRAS-dependent regulation of CCL2 secretion was also validated in our KRAS^G12V-ER^ pneumocyte cell line (Fig S4E). Furthermore, in vivo MRTX treatment of 3LL *Δ*NRAS tumour-bearing mice led to a significant downregulation of *Ccl2* expression in the tumour (Fig 4G), suggesting that tumour cells may be one of the main sources of this cytokine in the TME. To validate the role of KRAS-mediated regulation of CCL2 in the changes observed in the TME, we measured monocyte migration ex vivo. Results showed that migration was significantly abrogated when bone marrow-derived monocytes were cultured in conditioned medium from MRTX-treated cells and to a similar extent when cultured in medium from C*cl2*^-/-^ cells (Fig 4H). This data suggests that tumour cell specific KRAS^G12C^ inhibition, via inhibition of the secretion of CCL2, leads to an impaired recruitment of monocytes into the TME, which could constitute a mechanism that alleviates immunosuppression.

### KRAS^G12C^ inhibition in vivo increases T cell infiltration and activation

While a reduction in immunosuppressive populations in the TME constitutes a mechanism to improve anti-tumour immunity, immunological rejection can only be achieved by the specific activation of cytotoxic populations such as CD8+ T cells.

For the generation of an adaptive immune response, lymphocytes need to be primed by professional antigen presenting cells (APC), which in turn need to have been activated themselves by engulfment of tumour-specific antigens. Therefore, we sought to assess if the reduction of viability and increase of apoptosis caused by KRAS^G12C^ inhibition ((*14*) and Fig S1E) could affect dendritic cell (DC) activity in vitro. Indeed, the GFP+ Mutu DCs were able to phagocytose MRTX-treated CellTrace Violet (CTV) labelled 3LL *Δ*NRAS cells when co-cultured (Fig 5A), which constitutes the first step of DC activation. In addition, we found that in vivo MRTX-treatment increased the presence of APCs in the TME (Fig 5B), with a consistent increase in the expression of genes involved in antigen presentation (Fig 5C, S5A). Consistent with this finding, co-culture of tumour cells with DCs revealed that KRAS^G12C^ inhibition promoted the upregulation of activation markers MHCII and CD86 on DCs (Fig 5D). Interestingly MHC II upregulation seemed to be mediated by tumour cell secreted factors, while CD86 required the presence of tumour cells, suggesting that different mechanisms might be at play in the tumour cell-mediated activation of DCs (Fig S5B).

**Fig. 5.**
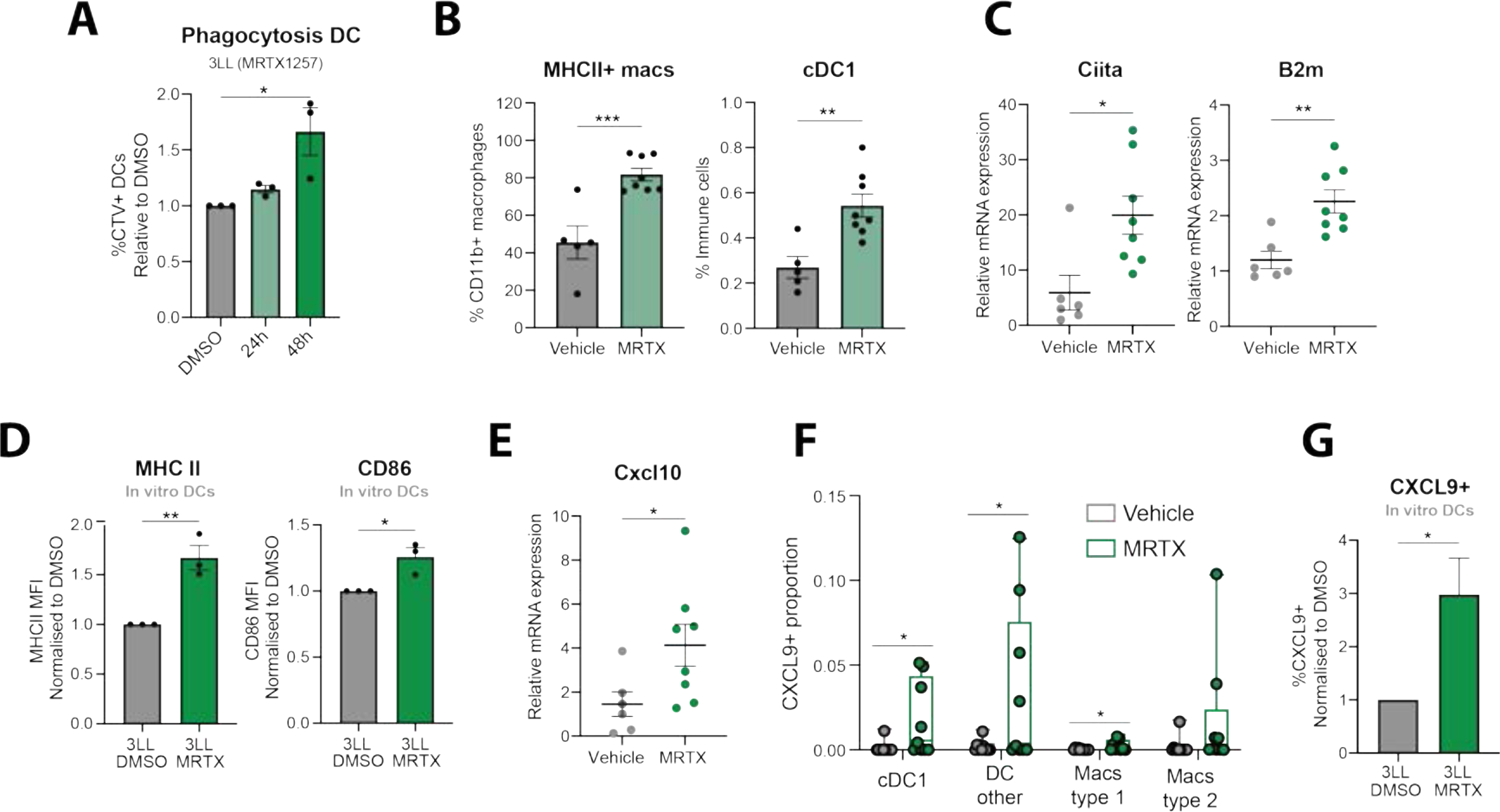
KRAS^G12C^ inhibition promotes APC activation. (A) Normalised percentage of GFP+ Mutu dendritic cells that have phagocytosed CTV+ 3LL *Δ*NRAS cells, previously treated with DMSO control, or 100nM MRTX1257 for 24h or 48h, measured by flow cytometry (n=3 independent experiments, One-way ANOVA, mean±SEM). (B) Flow cytometry analysis of 3LL *Δ*NRAS lung tumours treated with vehicle or 50mg/kg MRTX1257 for 7 days. Macrophages are gated as Live CD45+ CD11b+ CD24-CD64+ and cDC1s are obtained by Live, CD45+, CD11c+ CD24+ CD103+ gating (n=5 for vehicle n=8 mice for MRTX-treated, unpaired t test, mean±SEM). (C) qPCR data for 3LL *Δ*NRAS lung tumours treated as in (B) (2^-ΔΔCT^, unpaired t test, n=7 vehicle, n=8 treated, mean±SEM). (D) Normalised mean fluorescence intensity of MHCII and CD86 as measured by flow cytometry of DCs co-cultured with 3LL *Δ*NRAS cells previously treated with either DMSO or MRTX for 48h (pre-gated as GFP+, unpaired t test, n=3 independent experiments, mean±SEM). (E) qPCR data of Cxcl10 gene in 3LL *Δ*NRAS tumours, analyzed as in (C). (F) Proportion of CXCL9+ cells in each population, as detected by IMC, per ROI (n=11 vehicle, n=9 MRTX, unpaired t tests, mean±SEM). (G) Normalised percentage of CXCL9+ DCs after co-culture with 3LL *Δ*NRAS cells as in (D) (n=5 independent experiments, unpaired t test, mean+SEM).

Activated DCs are known to secrete CXCR3 ligands (CXCL9/10/11), a prominent feature of inflamed TMEs (*27*). We found that KRAS^G12C^ inhibition in vivo resulted in a higher expression of these T cell chemoattractants and their receptor (Fig 5E, Fig S5C). IMC analysis in the 3LL *Δ*NRAS lung tumours revealed that CXCL9 was mainly expressed in cells that also expressed markers for APCs (Fig S5D). Furthermore, while these tumours had negligible basal CXCL9 expression, CXCL9-expressing cells were mostly found among dendritic cells and macrophages in a subset of the MRTX treated tumours (Fig 5F). Mechanistically, we observed that the co-culture with MRTX-treated tumour cells was able to lead to the upregulation of CXCL9 in DCs in vitro (Fig 5G). This could not be recapitulated by conditioned medium incubation, suggesting that signals from MRTX-treated tumour cells, other than secreted factors, mediate this upregulation (Fig S5E).

Next, we evaluated whether the observed activation of APCs coincided with changes in the T cell compartment. We observed that the presence of all T cell compartments, particularly Foxp3+ Tregs was increased by MRTX in these lung tumours (Fig 6A). Consistent with the increased T cells, cytotoxicity genes were also significantly upregulated in MRTX-treated tumours (Fig 6B). Interestingly, NK cell infiltration was also increased in treated tumours (Fig S5F) which could be contributing to the increased expression of cytotoxicity genes. This increase in cytotoxicity was confirmed by the significantly increased presence of CD69+ and antigen-experienced (effector and memory) CD8+ T cells observed after MRTX treatment (Fig 6C and S5G).

**Fig. 6.**
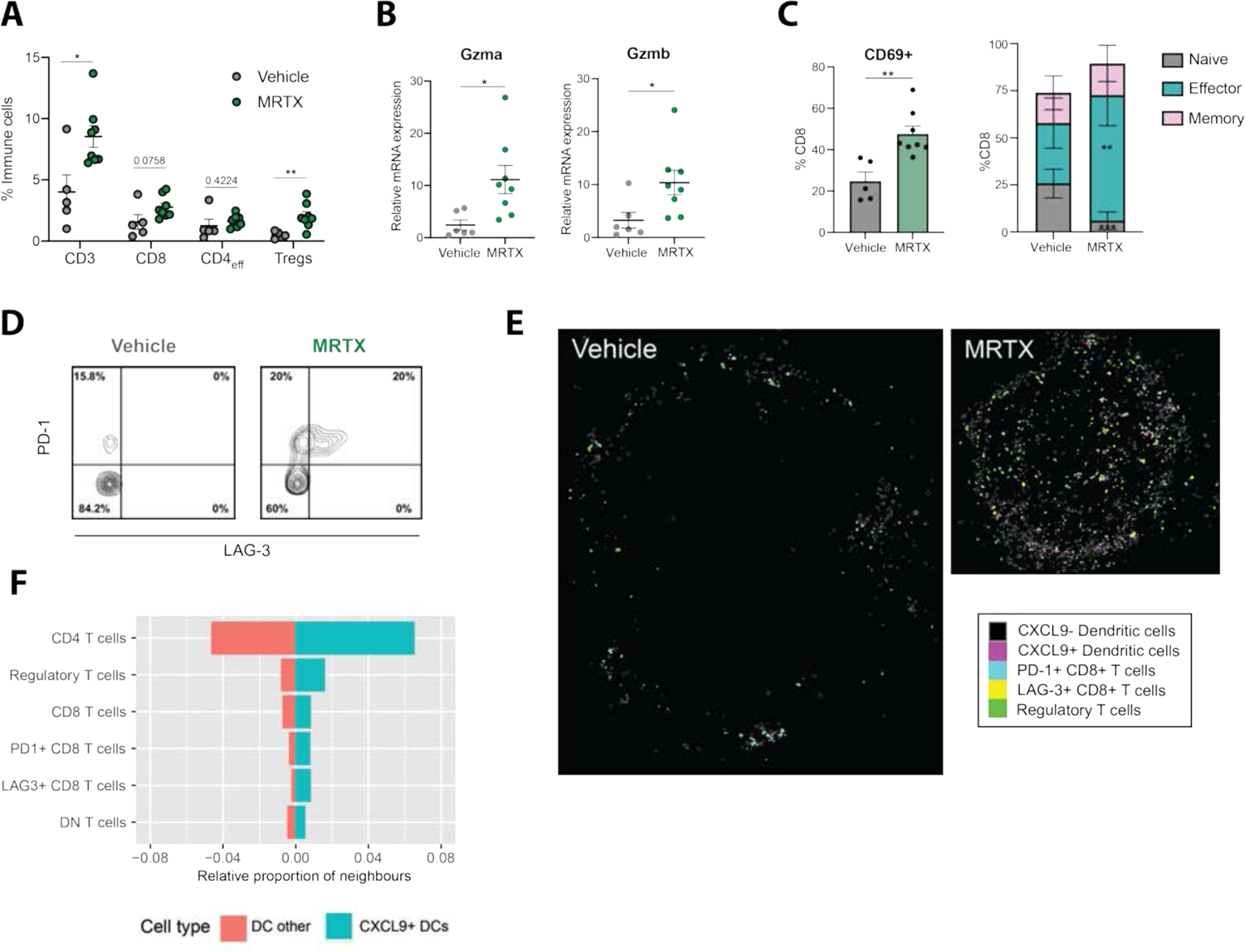
KRAS^G12C^ inhibition leads to T cell infiltration and activation. (A) Summary of T cell infiltration measured by flow cytometry in vehicle versus MRTX-treated lung tumours (n=5 for vehicle, n=8 mice for treated, unpaired t tests). (B) qPCR analysis of cytotoxicity genes in 3LL *Δ*NRAS lung tumour (2^-ΔΔCT^, unpaired t test, n=7 vehicle, n=8 treated, mean±SEM). (C) Flow cytometry analysis of CD8+ T cell phenotypes. Left: percentage of CD69+ CD8+ T cells in both treatment groups (n=5 vehicle, n=8 MRTX-treated, unpaired t test, mean±SEM). Right: Percentage of naïve (CD44-CD62L+), effector (CD44+ CD62L-) and memory (CD44+ CD62L+) CD8+ T cells, same analysis as on the left for each cell population. (D) Contour plot of PD-1 and LAG-3 expression on CD8+ cells in vehicle and MRTX-treated 3LL *Δ*NRAS lung tumour samples (graph shows one representative example for n=5 vehicle and n=8 MRTX treated samples). (E) Visualization of cell outlines as measured by IMC, of CXCL9 negative and positive DCs, PD-1+ and LAG-3+ CD8+ T cells and regulatory T cells in a vehicle and a MRTX-treated tumour. (F) Quantification of occurrence of the different T cell subsets in the neighbourhood of CXCL9+ and CXCL9-DCs, depicted as the average proportion of that cell type among all neighbors within 100px radius of the DCs subset.

Previous reports have shown that KRAS inhibition triggers an improved immune response that drives T cell exhaustion, resulting in sensitivity to immune checkpoint blockade (ICB) in immunogenic models of KRAS^G12C^-mutant cancer (*8, 19*). In our immune evasive 3LL *Δ*NRAS lung tumour model, we also observed via flow cytometry that PD1+ T cells were significantly increased after MRTX treatment (Fig S5H). A subset of these cells also expressed LAG-3 (Fig 6D) and we likewise found an upregulation of several other T cell exhaustion genes in our RNA-Seq analysis (Fig S5I).

As CXCL9 expression by DCs was previously described to be crucial to attract effector T cells (*27*), we further explored the relationship between the CXCL9+ DCs and the presence of different T cell subsets in the MRTX-treated tumours. There was a clear correlation between the abundance of CXCL9+ DCs and CD8+ T cells expressing PD1 and LAG-3 as well as Tregs (Fig S5J). Using the spatial information captured by IMC, we could also see that these cells are regularly found in close proximity to each other, with a clear enrichment of regulatory T cells and CD8+ T cells with an exhausted phenotype in the direct neighbourhood of CXCL9+ DCs compared to CXCL9-DCs (Fig 6E and 6F). Whether the CXCL9 expressing dendritic cells indeed play a role in recruiting these effector cells, or that activated T cells locally produce IFN*γ* that in turn induces the CXCL9 expression in the dendritic cells, cannot be deduced from this data.

Together, this data shows that tumour cell-specific KRAS^G12C^ inhibition in a mouse lung cancer model leads to a more inflamed TME, evidenced by an activation of APCs and a strong increase in the presence of activated T cells that could exert cytotoxic actions on the tumour cells, but also display an exhausted phenotype.

### KRAS^G12C^ inhibition synergizes with checkpoint blockade only in intrinsically immunogenic tumours

An increased presence of exhausted T cells and augmented IFN responses suggest that MRTX treatment has the potential to sensitize these tumours to ICB. Nevertheless, in this immune-resistant model (*28*), addition of an anti-PD1 antibody (Fig 7A) or a combination of anti-PD-L1 and anti-LAG3 antibodies (Fig S6A) did not improve the response to KRAS^G12C^ inhibition alone, nor did it enhance the TME remodeling driven by KRAS^G12C^ inhibition (Fig 7B). We found that the lack of therapeutic response observed was not due to insufficient antigen presentation by the tumour cells, as re-expression of the epigenetically silenced H2Kb by treatment with the DNA methyltransferase inhibitor Decitabine in these cells did not improve responses to MRTX+PD1 (Fig S6B and S6C).

**Fig. 7.**
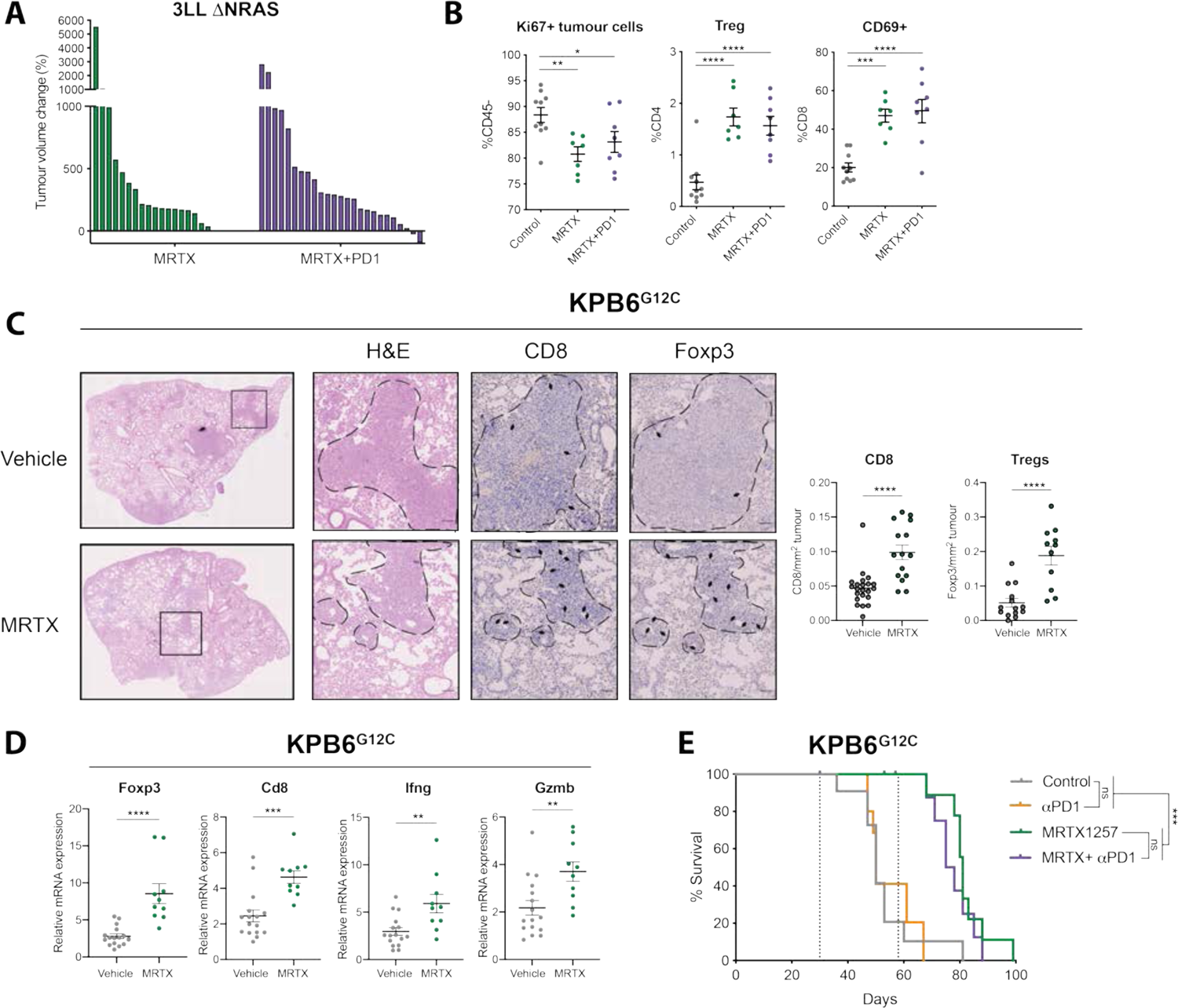
KRAS^G12C^ inhibition does not synergize with ICB in immune refractory tumours. (A) Tumour volume change after two weeks of treatment of 3LL *Δ*NRAS-tumour bearing mice with either 50mg/kg MRTX1257 only (n=9 mice) or MRTX1257 and anti-PD1 (n=10 mice). Each bar represents a tumour. (B) Flow cytometry analysis of proliferating tumour cells (CD45-Ki67+), Tregs (CD3+ CD4+ Foxp3+) and activated T cells (CD69+ CD8+) in vehicle (n=10), MRTX (n=7) or MRTX plus anti-PD1 (n=8) treated (two week treatment) 3LL *Δ*NRAS lung tumours (One way ANOVA, mean±SEM). (C) Immunohistochemistry analysis and quantification for CD8 (n=4 mice per group) and Foxp3 (n=3 mice per group) in KPB6^G12C^-tumour bearing lungs after 7 days of vehicle or 50mg/kg MRTX1257 treatment (each dot represents one tumour, unpaired t test, mean±SEM). (D) qPCR analysis of immune genes in vehicle (n=15 tumours) or MRTX-treated (n=10 tumours) KPB6^G12C^ lung tumours (2^-ΔΔCT^, Unpaired t test, mean±SEM). (E) Survival of KPB6^G12C^lung tumour-bearing mice treated with vehicle (+IgG control, n=6 mice), MRTX1257 (+IgG control, n=6 mice), anti-PD1 (n=5 mice) or combination (n=4 mice, Log-Rank Mantel Cox test).

To extend our findings, we made use of the KPB6^G12C^ cell line, which has been established from the KRAS^LSL_G12D/+^;Trp53^fl/fl^ mice (KP) and genetically engineered to express a KRAS^G12C^ mutation (*25*). Due to the very low number of clonal somatic SNVs, this model develops immune cold lung tumours (*25*). Orthotopic KPB6^G12C^ lung tumours were highly sensitive to KRAS^G12C^ inhibition (Fig S6D). Treatment of KPB6^G12C^ lung tumour-bearing mice with MRTX for a week led to increased T cell infiltration into the tumours (Fig 7C) accompanied by an upregulation of immune genes (Fig 7D). Similar to our findings in 3LL *Δ*NRAS, we found a significant increase in CD8+ T cell and Tregs and increased *Ifng* and *Gzmb* expression, suggesting increased cytotoxicity. However, in this alternative immune resistant model, no synergism occurred between KRAS^G12C^ inhibition and ICB, and mice in all treatment groups succumbed to disease (Fig 7E). We therefore concluded that despite the profound TME remodeling triggered by KRAS^G12C^ inhibition, it may not be sufficient to render highly immune resistant tumours sensitive to ICB.

We therefore investigated the effects of KRAS^G12C^ inhibition in a new immunogenic model of KRAS-mutant lung cancer. The KPAR^G12C^ cell line has been shown to be immunogenic as its growth is impaired by the adaptive immune system (*25*). While sensitivity to KRAS^G12C^ inhibition in vitro was reduced compared to the 3LL *Δ*NRAS cell line (Fig S7A), the responses seen in vivo were much stronger, with most lung tumours shrinking more than 75% (Fig 8A). Treatment of subcutaneous KPAR^G12C^-tumour bearing mice with a KRAS^G12C^ inhibitor also resulted in outstanding tumour control, with two out of seven mice achieving complete responses (Fig 8B). These responders were resistant to tumour re-challenge, suggesting the development of immune memory. In contrast, the responses of 3LL *Δ*NRAS tumour-bearing mice to KRAS^G12C^ inhibition were not supported by the adaptive immune system. There were no long-term responses, with all mice relapsing on treatment, and responses were comparable in immunocompetent and immunodeficient mice (Fig S7B).

**Fig. 8.**
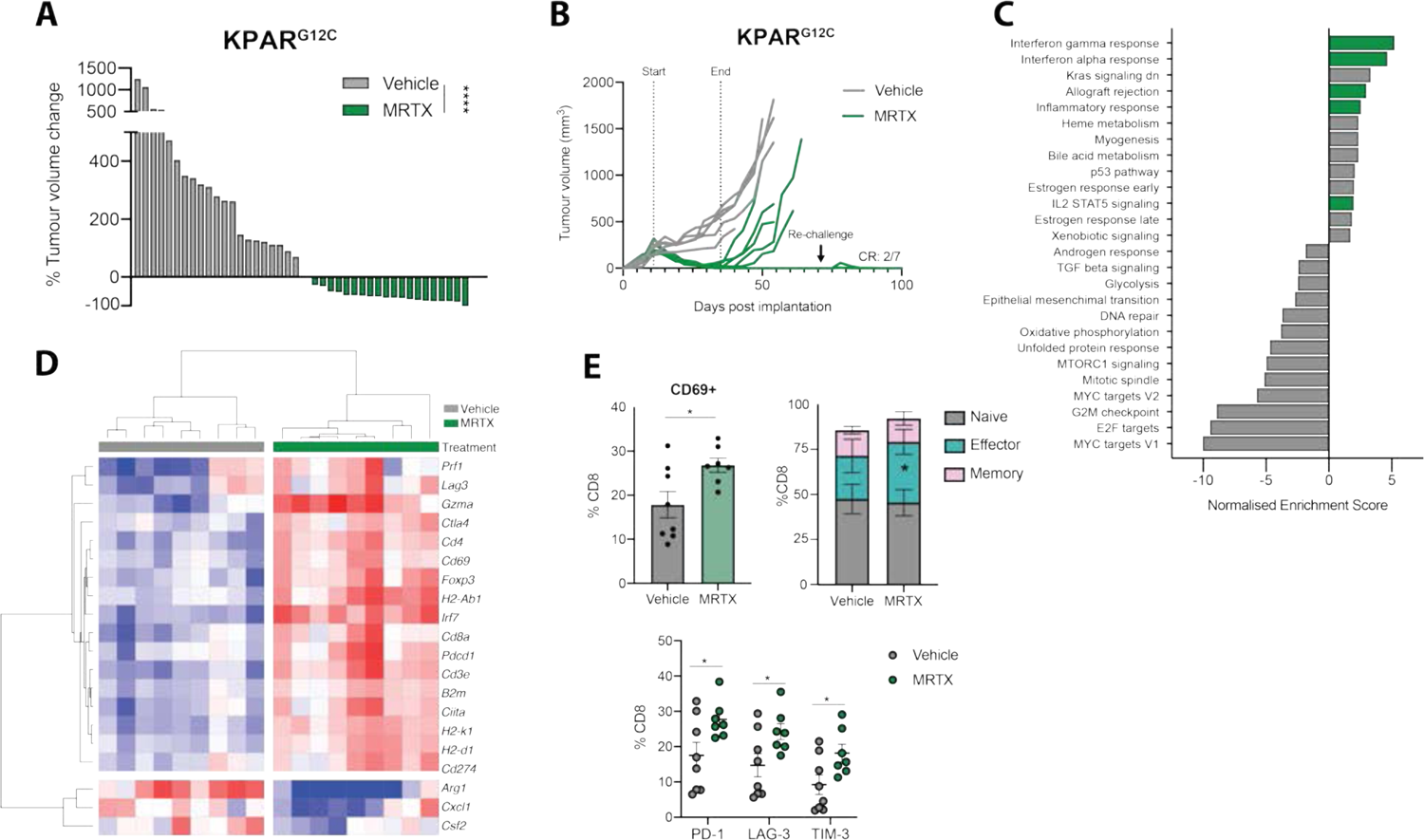
In an immunogenic model, MRTX-driven immune responses drive complete tumour rejection. (A) Tumour volume change after seven days of treatment of KPAR^G12C^-tumour bearing mice with either vehicle (n=3 mice) or MRTX849 (n=2 mice). Each bar represents a tumour, Mann-Whitney test. (B) Growth of subcutaneously implanted KPAR^G12C^-tumours treated with either vehicle or 50mg/kg of MRTX849 for two weeks. At day 71, remaining mice were re-challenged with KPAR^G12C^ cells in the opposite flank, which did not give rise to tumours. (C) Summary of significantly (FDR q<0.05) up- and down-regulated pathways in MRTX849 (50mg/kg, 6 days) versus vehicle-treated KPAR^G12C^ lung tumours (MSigDB Hallmarks), n=9 tumours per group (3 mice). (D) Heatmap showing mRNA expression from RNA-Seq of KPAR^G12C^ tumours treated for 6 days with 50mg/kg MRTX849. Gene expression is scaled across all tumours. (E) Flow cytometry analysis of KPAR^G12C^-bearing lungs treated with either vehicle (n=8 mice) or 50mg/kg MRTX849 (n=7 mice) for 6 days, showing increased CD69+ CD8+ T cells (top left), increased CD44+CD62L-effector CD8+ T cells (top right) and increased checkpoint molecule expression on CD8+ T cells (below) after KRAS inhibition (all statistics are Student’s t tests, mean±SEM).

We then assessed the effects of KRAS^G12C^ inhibition on the remodeling of the TME in KPAR^G12C^ lung tumours. RNA-Seq from MRTX-treated tumours showed an upregulation of immune-related gene sets, confirming our observations in the 3LL *Δ*NRAS and KPB6^G12C^ models (Fig 8C). Genes encoding for T cell infiltration (*Cd3e*, *Cd4*, *Cd8a*, *Foxp3*), T cell activation (*Prf1, Cd69, Gzma, Pdcd1, Ctla4, Lag3*), IFN responses (*Irf7, Irf9, Cd274*) and antigen presentation (*H2-Ab1, H2-K1, H2-D1, Ciita, B2m*) were upregulated after treatment while immunosuppressive cytokines (*Cxcl1, Csf2*) and markers of tumour-promoting myeloid populations (*Arg1*) were downregulated (Fig 8D and S7C). Flow cytometric analysis revealed a significant upregulation of activated, antigen-experienced and exhausted T cells (Fig 8E) and NK cells (Fig S7D) as well as a remodelling of the myeloid compartment, with a reduction of neutrophils and increased APC activation (Fig S7D) similar to previous models examined.

In this immunogenic model, where early treatment with anti-PD1 alone confers therapeutic benefit (Fig 9A), treatment of orthotopic tumour-bearing mice with MRTX alone led to complete responses in 28% of the mice (Fig 9B). Furthermore, the percentage of complete responders was improved (66%) when KRAS^G12C^ inhibition was administered together with anti-PD1 immunotherapy treatment, even while treatment started later at a time point when single anti-PD1 therapy was no longer effective (Fig 9B). The synergy between KRAS^G12C^ inhibition and anti-PD1 in this model was also reflected by the composition of the TME, with a further increase in immune infiltration and activation genes observed in the combination treatment (Fig S7E).

**Fig. 9.**
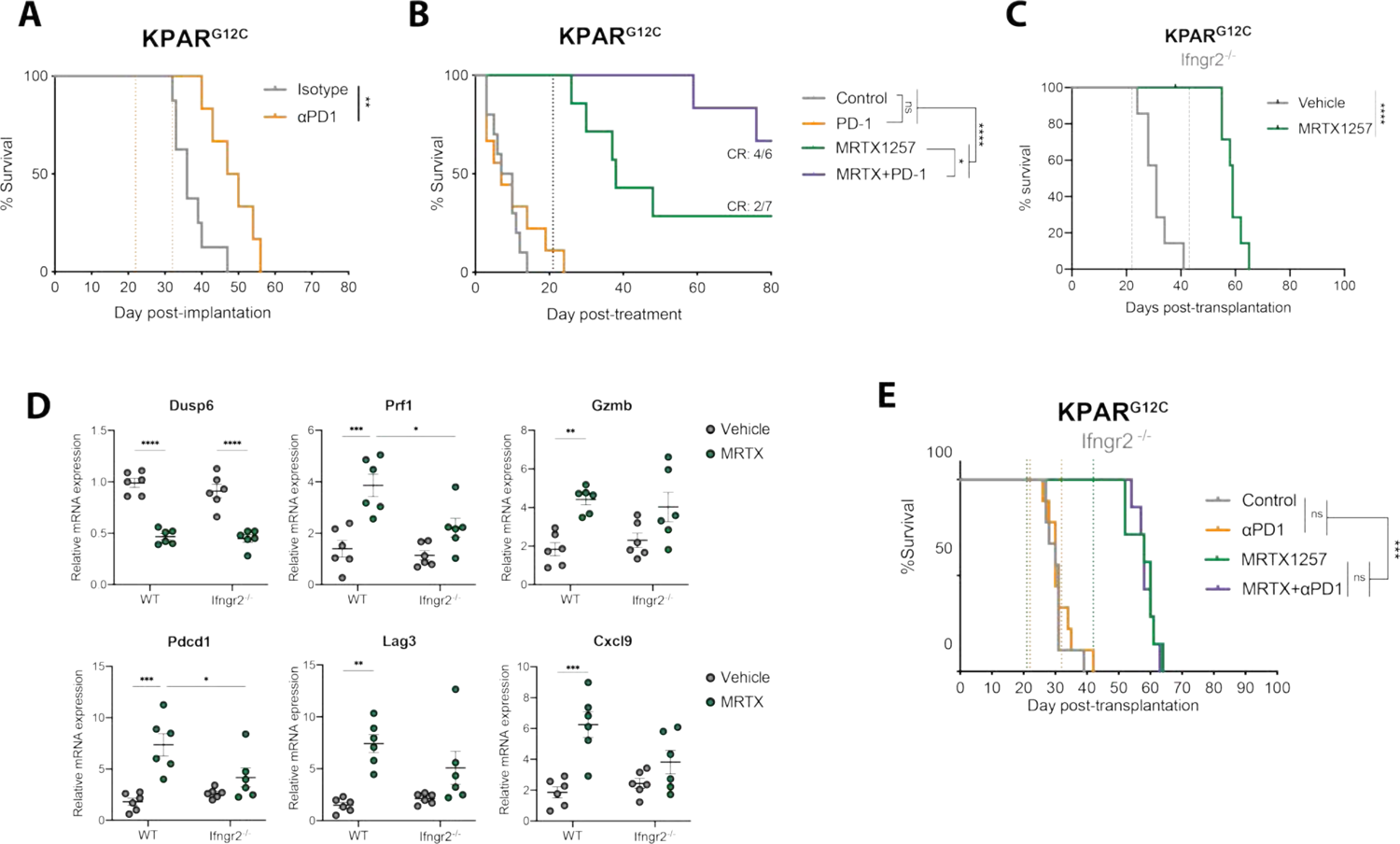
Synergy with anti-PD-1 requires an intact tumor cell-intrinsic IFN response. (A) Survival of KPAR^G12C^ lung tumour-bearing mice after treatment with IgG control (n=8 mice) or 10mg/kg anti-PD-1 (n=6 mice). Dotted lines represent start and end of treatment, respectively, Log-Rank Mantel Cox test. (B) Survival of KPAR^G12C^ lung tumour-bearing mice after treatment with Vehicle (+IgG control, n=6 mice), 10mg/kg anti-PD-1 (n=8 mice), 50mg/kg MRTX1257 (+ IgG control, n=4 mice) or both (n=6 mice). Dotted line represents end of treatment, Log-Rank Mantel Cox test. (C) Survival of KPAR^G12C^ *Ifngr2*^-/-^ lung tumour-bearing mice after treatment with vehicle or 50mg/kg MRTX1257 (n=7 mice per group). Dotted lines represent treatment start and end, respectively, Log-Rank Mantel Cox test. (D) qPCR analysis of KPAR^G12C^ WT or *Ifngr2*^-/-^ lung tumours treated with vehicle or MRTX1257 for 4 days (n=6 tumours per group, mean±SEM, 2^-ΔΔCt^). Each dot represents a lung tumour, one way ANOVA. (E) Survival of KPAR^G12C^ Ifngr2^-/-^ lung tumour-bearing mice after treatment with Vehicle (+IgG control, n=9 mice), 10mg/kg anti-PD-1 (n=9 mice), 50mg/kg MRTX1257 (+ IgG control, n=7 mice) or both (n=7 mice). Dotted line represents start and end of treatment for MRTX (green) and anti-PD-1 (orange), Log-Rank Mantel Cox test.

Using the KPAR^G12C^ cell line, we validated the KRAS-dependent regulation of IFN responses, driven through MYC (S7F), highlighting the universality of this novel mechanism. We then made use of the immunogenicity of this model and examined the role of tumour cell-intrinsic IFN*γ* signaling in the long-term therapeutic effect of KRAS^G12C^ inhibitors. To this end, we generated *Ifngr2*^-/-^ KPAR^G12C^ cells (Fig S7G), which are insensitive to IFN*γ*, while the KRAS inhibitor driven upregulation of IFN genes remains unaffected (Fig S7H and S7I). We observed that complete responses to KRAS^G12C^ inhibition in vivo were dependent on tumour cell-intrinsic IFN signaling, as all mice bearing tumours formed by *Ifngr2*^-/-^ KPAR^G12C^ cells relapsed after MRTX treatment (Fig 9C), while their sensitivity to KRAS inhibition in vitro remained unaffected (Fig S7J). Similarly, KRAS target gene *Dusp6* reduction in vivo was comparable in KPAR^G12C^ WT and *Ifngr2*^-/-^ tumours (Fig 9D, top left). On the contrary, we observed that in *Ifngr2*^-/-^ tumours, the increase in T cell cytotoxicity (*Prf1*, *Gzmb*), activation (*Pdcd1, Lag3*) and myeloid cell activation (*Cxcl9*) in response to KRAS^G12C^ inhibition was significantly attenuated, probably contributing to the decreased long-term therapeutic efficacy of the inhibitor in this model. Furthermore, the synergism between MRTX and anti-PD1 treatment observed in this model was completely abrogated in *Ifngr2^-/-^* tumours (Fig 9E).

Together, these data suggest that in an immunogenic tumour, KRAS^G12C^ inhibition can stimulate anti-tumour immunity, drive complete tumour rejection in a subset of mice and sensitize tumours to ICB, resulting in increased complete responses when both treatments are combined. In addition, this process is dependent on the tumour cell-intrinsic ability to respond to IFN*γ*, which is regulated by KRAS signaling and contributes to long-term therapeutic efficacy of KRAS^G12C^ inhibition.

## DISCUSSION

KRAS^G12C^ inhibitors have shown promising clinical activity in KRAS^G12C^-mutant NSCLC patients (*9, 10*). However, similarly to other targeted therapies, early clinical results indicate that drug resistance frequently arises, resulting in clinical relapses (*11, 12*). KRAS^G12C^ inhibitors not only affect the survival of cancer cells but can also mediate immunomodulatory effects by reversing KRAS-driven immunosuppressive mechanisms and generate a TME that is more favorable for an anti-tumour immune response (*8, 19, 21*). This knowledge has served as a rationale to investigate clinical combinations of KRAS^G12C^ inhibitors with anti-PD1 or PD-L1 antibodies (*29*). However, previous studies have only validated this combination using mouse models that are highly ICB-responsive (*8, 19*). Here we show that despite the profound TME remodeling caused by KRAS^G12C^ inhibition, this drug combination may not be sufficient to elicit durable responses in tumour models that are intrinsically resistant to immune checkpoint blockade.

Understanding the mechanism of action of KRAS^G12C^ inhibitors and how they can modulate the TME may lead to the identification of additional combination strategies for those patients that will not benefit from the dual inhibition of KRAS^G12C^ and PD1. Our analysis has gained insight into the different mechanisms by which oncogenic KRAS signaling mediates immune evasion in lung cancer. It has already previously been described that oncogenic KRAS regulates the expression of cytokines and chemokines that can modulate the TME (*22, 30, 31*). Here we show that KRAS^G12C^ inhibition reduces the secretion of monocyte and neutrophil chemoattractants by the tumour cells, which results in an impaired infiltration of these immune suppressive cell types in the TME. Reprograming myeloid populations by targeting selected cytokines or their receptors, like CCR2 (*32*) or CXCR2 (*33*) has been proposed as a mechanism to enhance response to immunotherapies (*34*). Treatment with KRAS^G12C^ inhibitors leads to modulation of various C(X)CL ligands secreted by tumours cells and can thus indirectly reduce immunosuppressive populations without associated toxicities. However, the identity of the KRAS regulated cytokines appears to vary between tumour types.

Another mechanism by which oncogenic KRAS drives immune evasion is by inhibiting IFN responses. We have shown that KRAS^G12C^ inhibitor treatment releases the inhibition of IFN signaling pathway genes in all the models that we have analyzed. Moreover, activation of oncogenic KRAS in type II pneumocytes inhibits IFN pathway expression, suggesting that this is a conserved mechanism in the lung. Mechanistically, KRAS inhibits IFN gene expression via regulation of the oncogene MYC, which is consistent with previous observations in pancreatic cancer (*26*). Importantly, KRAS^G12C^ inhibition enhances tumour cell sensitivity to type I and II interferons and results in an increased IFN pathway activation in vivo. This is especially important as IFN responses are crucial for anti-tumour immunity and clinical responses to immunotherapies (*23, 24, 35, 36*). By knocking out the IFN*γ* receptor in tumour cells, we have demonstrated that tumour cell intrinsic IFN signaling is necessary to achieve long-lasting therapeutic responses to KRAS^G12C^ inhibitors in vivo. We have therefore expanded beyond the known role of IFN signaling in the response to immune therapies (*37, 38*), showing that an intact interferon response is also required for durable immune responses to a targeted therapy such as KRAS^G12C^ inhibition.

As a consequence of the KRAS-dependent regulation of IFN responses, treatment with KRAS^G12C^ inhibitors increases antigen presentation. Oncogenic KRAS has previously been linked to reduced expression of MHC class I molecules (*39, 40*). Reversion of this immune evasion mechanism can boost T cell recognition rendering tumour cells more susceptible to immune cell attack. Additionally, the cell death induced by KRAS^G12C^ inhibitors could also trigger an adaptive T cell response due to the release of dead cell-associated antigens.

Consistent with this, we observe both in vitro and in vivo that KRAS^G12C^ inhibition indirectly increases professional antigen presentation by promoting the activation of APCs accompanied by an increase of CXCR3-binding chemokine expression by DCs. These effects of KRAS^G12C^ inhibition can explain the elevated CD8+ T cell recruitment and the increased T cell activation that we observe upon treatment. Importantly, these characteristics are a prominent feature of ‘inflamed’ TMEs (*27, 41*), which are more likely to respond to immunotherapy.

KRAS^G12C^ inhibition alleviates immunosuppressive mechanisms and enhances the infiltration and activation of cytotoxic T cells, accompanied by an increase in checkpoint molecule expression, such as PD1 and LAG-3, even in a very immunosuppressive model like the 3LL *Δ*NRAS tumours. This TME could be considered optimal for the addition of immune checkpoint blockade inhibitors to potentiate a T-cell dependent immune response (*42*). However, the combination of KRAS^G12C^ inhibition with anti-PD1 was only synergistic in the immunogenic tumour model (KPAR1.3 G12C), but not in the two models that were intrinsically resistant to ICB, one ‘cold’ tumour model lacking neoantigens (KPB6^G12C^) and one ‘T cell excluded’ model which evades anti-tumour immunity by downregulating MHC and recruiting immunosuppressive myeloid cells (3LL *Δ*NRAS). While we cannot rule out a beneficial effect of the combination in all tumours with immune refractory TMEs, as our models certainly do not cover the whole spectrum of immunogenicity observed in NSCLC patients, it will be of utmost importance to identify which patients can benefit from the addition of anti-PD1 inhibitors to KRAS^G12C^ inhibitors and to investigate additional therapeutic strategies for the remaining patients. Our mouse models offer the opportunity for future investigation on additional combinatorial therapies as the therapeutic approach could differ depending on the mechanism of immune evasion. ‘Cold’ tumour-bearing patients may benefit from the addition of drugs aimed to increase antigen load, such as chemotherapy, radiotherapy, epigenetic modulators or STING agonists (*43*), whereas targeting immunosuppressive cells could be a valid therapeutic strategy for ‘T cell excluded’ tumours. KRAS^G12C^ inhibitors can already decrease some myeloid immune suppressive populations, however treatment consistently results in an increase in the infiltration of Tregs, which inhibit cytotoxic T cell activity and might represent an alternative target for combination therapy (*44*).

Several preclinical studies, including this one, have demonstrated that combinations of KRAS^G12C^ inhibitors with anti-PD1 can clearly result in therapeutic benefit in immunogenic mouse cancer models (*8, 19*). Based on these data, a number of different clinical trials are underway testing combinations of KRAS^G12C^ inhibitors and PD1 pathway immune checkpoint blockade, such as KRYSTAL-1, KRYSTAL-7, CodeBreak 100 and CodeBreak 101, with results eagerly awaited. With these and other clinical trials already running, there are still open questions that need to be addressed in order to set up the basis for patient stratification. Our findings are particularly relevant for those patients with highly immune refractory TMEs as they could benefit instead from other combination strategies. While it is likely that the ongoing trials of combinations of KRAS^G12C^ inhibitors with immunotherapies will be beneficial for a subset of KRAS-mutant NSCLC patients, however, this study has highlighted the need for additional treatment strategies in highly immune refractory patients. In particular, it should be noted that most of these combination clinical trials, with the exception of KRYSTAL-7, do not exclude prior treatment with immunotherapy, and are therefore likely to be enriched with patients whose tumours show either intrinsic or acquired resistance to immune checkpoint blockade. Extrapolating from the preclinical studies reported here, such patients may be less likely to benefit from combinations of KRAS^G12C^ inhibitors with immunotherapies.

While KRAS^G12C^ inhibitors have only recently been approved for clinical use, MEK inhibitors, targeting the MAPK pathway downstream of KRAS, have been used for some time and can result in similar tumour cell-intrinsic immunomodulatory changes (*45*) and in some cases have shown to ameliorate anti-tumour immunity (*46–49*). However, the positive effects in the TME caused by the tumour cell-intrinsic changes can be reduced by the detrimental effects of MEK inhibition on immune cells (*8*). Moreover, although combinations of inhibitors targeting MAPK pathway plus anti-PD1 can improve clinical outcomes, they do so at the expense of increased toxicities (*50, 51*). In contrast, KRAS^G12C^ inhibitors offer the unique ability to improve anti-tumour immunity via a myriad of mechanisms discussed above while not affecting MAPK signaling in non-tumour cells, including those involved in the anti-tumour immune response.

Consequently, unlike other targeted therapies that do not specifically target oncogenic mutant proteins, KRAS^G12C^ inhibitors have the potential to achieve long-term survival which is dependent on the activation of anti-tumour immune responses in immunogenic tumours. The tumour cell-specific activity of KRAS^G12C^ inhibitors provides an unprecedented opportunity to investigate combinations of multiple therapeutic approaches without producing excessive toxicity profiles. Several clinical trials are testing combinations of KRAS^G12C^ inhibitors with other targeted therapies, including MEK inhibitors (*29*). It will be important to validate that the beneficial effects upon the TME are not lost when these two drug classes are combined. With that in mind, it is possible the “vertical” combinations of KRAS^G12C^ inhibitors with SHP2 inhibitors upstream or CDK4/6 inhibitors downstream may be more promising, as both these drug types have also been shown to produce positive immunomodulatory effects (*20, 52, 53*).

## MATERIALS AND METHODS

### Study design

The objective of this study was to examine non-tumour cell intrinsic effects of KRAS^G12C^ inhibitors. We performed controlled (non-blinded) laboratory experiments using cancer cell lines to examine the effects of KRAS^G12C^ inhibitors on gene and protein expression and co-culture systems with immune cells to assess indirect effects of the drug treatment on different cell populations. For all in vitro experiments a minimum of two biological replicates (independent experiments) were acquired.

We also used transplantable murine lung cancer models to assess the effects of KRAS^G12C^ inhibitors in non-blinded randomized studies (alone or in combination with ICB) on mouse survival. Endpoints were pre-defined and not modified throughout the duration of the study and mice whose cause of death could not be attributed to lung tumours were excluded. Other in vivo experiments aimed to investigate the TME, by combining RNA, flow cytometry and imaging mass cytometry data. Sample size was chosen empirically based on results of previous studies and no datapoints, including outliers, were excluded from these analyses.

### In vivo tumour studies

All studies were performed under a UK Home Office approved project license and in accordance with institutional welfare guidelines.

For subcutaneous tumour injection, cells were mixed 1:1 with GeltrexTM matrix (ThermoFisher) and 400,000 3LL *Δ*NRAS or 150,000 KPAR^G12C^ cells were injected in a total volume of 100μl subcutaneously into one flank of 8-week-old C57BL/6 mice. Tumour growth was followed twice a week by caliper measurements and tumours were left to grow not larger than 1.5cm in diameter following a UK Home Office approved project license.

For orthotopic growth, 10^6^ 3LL *Δ*NRAS or 150,000 KPAR^G12C^ cells were injected in PBS in a total volume of 100μl in the tail vein of 8-week-old C57BL/6 mice. Mouse weight was monitored regularly as a measure of tumour growth and mice were sacrificed if weight loss was over 15% as per the UK Home Office approved project license. Tumour burden was also assessed by regular Computed Tomography (CT) scanning of the lungs. Briefly, mice were anesthetized by inhalation of isoflurane and scanned using the Quantum GX2 micro-CT imaging system (Perkin Elmer) at a 50μm isotropic pixel size. Serial lung images were reconstructed and tumour volumes subsequently analyzed using Analyse (AnalyzeDirect).

For therapeutic experiments, mice were treated daily via oral gavage with 50mg/kg MRTX1257 (Mirati Therapeutics), 50mg/kg MRTX849 (MedChemExpress) or 10% Captisol® (Ligand) in 50mM citrate buffer (pH 5.0) as vehicle control.

For ICB treatments, mice were administered 10mg/kg anti-PD1 (clone RMP1-14, BioXcell), 10mg/kg anti-PD-L1 (clone 10F.9G2, BioXCell) and/or 10mg/kg anti-LAG3 (clone C9B7W, BioXCell) or isotype control (10mg/kg IgG2b and 5mg/kg Syrian hamster IgG2) dissolved in PBS at a dose of 4μl/g mouse intraperitoneally twice a week for a total of four doses.

### Cell lines

NCI-H23, NCI-H358 were obtained from the Francis Crick Institute Cell Services Facility. 3LL *Δ*NRAS were generated as previously described (*14*). KRAS^G12V^-ER pneumocytes were generated as previously described (*14*). KPAR-KRAS^G12C^ and KPB6-KRAS^G12C^ were generated as previously described (*25*). CT26-KRAS^G12C^ were obtained from Mirati Therapeutics (Briere). MutuDC cells were kindly provided by Dr. Caetano Reis e Sousa. KPAR-KRAS^G12C^ were maintained in DMEM and MutuDC in IMDM. The rest of the cell lines were cultured in RPMI. Medium was supplemented with supplemented with 10% fetal calf serum, 4mM L-glutamine (Sigma), 100units/ml penicillin and 100mg/ml streptomycin (Sigma). Cell lines were tested for mycoplasma and were authenticated by short-tandem repeat (STR) DNA profiling by the Francis Crick Institute Cell Services facility. Cells were allowed to grow for not more than 20 sub-culture passages.

### In vitro drug treatments

Cells were plated at an appropriate density and left to grow for at least 24h before drug treatment. Drugs were administered in fresh medium and samples were collected indicated time points for downstream analysis. Trametinib (10nM), GDC0941 (500nM), Everolimus (100nM), Ruxolitinib (500nM) and Decitabine (250nM) were obtained from Selleckchem. IFNAR blocking antibody (20mg/ml) was obtained from BioXcell. 4-OHT (500nM) was obtained from Sigma-Aldrich. ARS-1620 (2mM) was a generous gift from Araxes Pharma, LLC. MRTX1257 (100nM) was a generous gift from Mirati Therapeutics. Unless otherwise stated, concentrations used for in vitro experiments are indicated in brackets. Human and mouse recombinant IFN*α*/*β*/*γ*(all from Biolegend) were used at a concentration of 100ng/ml.

### In vitro viability assay

For viability assays, the CellTiter-Blue assay (Promega) was used. Cells were grown in 96-well plates and treated appropriately for 72h. At the end of the experiment, 5μl of the CellTiter-Blue reagent was added to each well and the reaction was incubated for 90 minutes in the incubator at 37°C. Fluorescence was subsequently measured using and EnVision plate reader (Perkin Elmer) with excitation/emission wavelengths of 560/590nm.

### Immunoblotting

Cells were lysed using 10X Cell Lysis Buffer (Cell Signaling Technologies), supplemented with EDTA-free protease inhibitor cocktail tablets (Roche), 1mM PMSF and 25mM NaF. 15-20mg of protein was diluted in NuPAGE LDS Sample Buffer (4X, Thermo Fisher) and samples were loaded onto NuPAGE 4-12% Bis-Tris protein gels (Thermo Fisher). Protein transfer to PVDF membranes was performed using the Trans-Blot Turbo Transfer System (BioRad) or standard manual transferring techniques. For antibody detection, horseradish peroxidase-conjugated antibodies were used (GE Healthcare) and data was developed using an Amersham Imager 600 (GE Healthcare) or standard film techniques. Immunoblot quantification was performed using ImageJ software (NIH).

Antibodies directed against phospho-ERK (T202/Y204, #9101), ERK (#9107), phospho-AKT (S473, #9271), AKT (#2920), phospho-S6 (S235/236, #2211), S6 (#2317), phosphor-STAT1 (T701, #9167), STAT1 (#9172), STAT2 (#4594) were obtained from Cell Signaling Technologies (CST). Pan-RAS antibody was obtained from Merck Millipore (MABS195), Vinculin (V9131) from Sigma-Aldrich and c-MYC (ab39688) from Abcam.

### RAS pulldown assay

Active Ras was measured using the Ras Activation Assay Kit from Millipore following manufacturer’s instructions. Briefly, cells were lysed in Mg^2+^ Lysis Buffer (MLB, 5% NP40, 750mM NaCl, 125mM Hepes, 50mM MgCl2, 5mM EDTA, 10% Glycerol) containing protease inhibitors. 500μg of protein was incubated with RAF-RBD containing agarose beads and rotated for 75 min at 4°C. Pulled down protein was then analysed by Immunobloting, using 20μg of non-bead incubated protein to normalise for total Ras levels.

### CRISPR-Cas9 knockout

Phosphorylated and annealed *Ccl2*-targeting (sgRNA 1 3’-gRNA-‘5: ACACGTGGATGTCTCCAGCCG and sgRNA 2: (5’-gRNA-‘3): GCAAGATGATCCCAATGAGT) or Ifngr2-targeting sgRNAs (3’-gRNA-‘5: AGGGAACCTCACTTCCAAGT) were cloned into target vector px458-pSpCas9(BB)-2A-GFP (Addgene #48138) or px459-pSpCas9(BB)-2A-Puro (Addgene #62988), respectively. 3LL *Δ*NRAS or KPAR^G12C^ cells were transfected using Lipofectamine (Thermo Fisher) with the px458 vector and FACS sorted for GFP expression or selected using puromycin treatment. Cells were then single cell cloned before KO screening via Sanger Sequencing and protein analysis via ELISA or FACS.

### siRNA transfection

siGENOME siRNAs against mouse Stat1, Stat2 or Myc (Dharmacon) were transfected at a final concentration of 50nM using DharmaFECT 4 transfection reagent (Dharmacon). The transfection complex was incubated for 20-40 minutes before adding dropwise to freshly seeded cells. As a control, cells were either Mock-transfected (no siRNA) or transfected with a siGENOME RISC-free control (Dharmacon).

### Quantitative RT-PCR

RNA was extracted using RNeasy Mini Kit (QIAGEN) following manufacturer’s instructions. For *in vivo* tumour samples, tumours were individually isolated from the lungs, lysed and homogenised using the QIAshredder (QIAGEN) following manufacturer’s instructions prior to RNA extraction. SuperScript II Reverse Transcriptase (Thermo Fisher) was then used to generate cDNA.

qPCR was performed using SYBR Green FAST Master Mix (Applied Biosystems). For a list of primers used see table 1. Gene expression changes relative to the housekeeping genes were calculated using the ΔΔCT method.

**Table 1.**
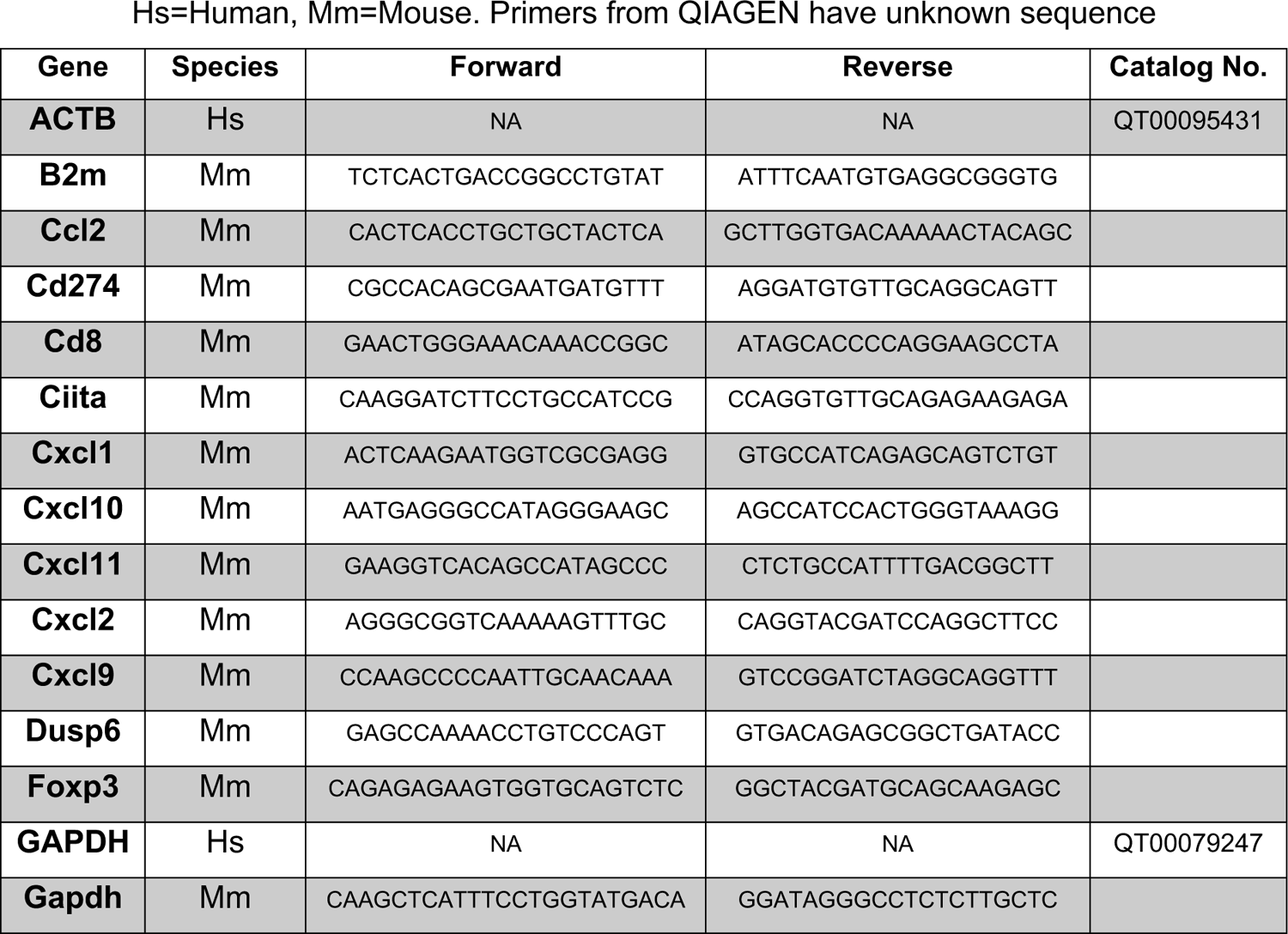
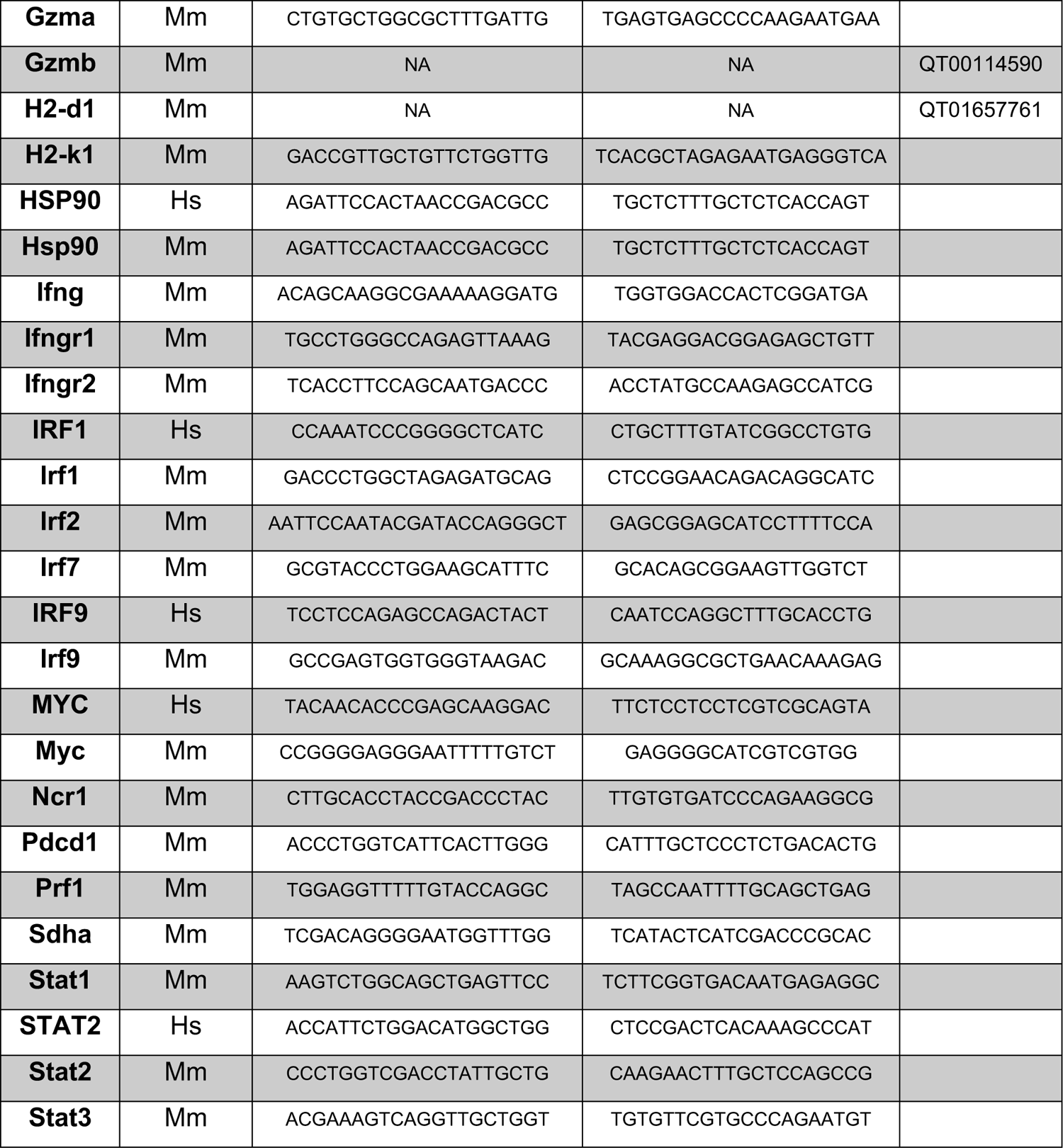
List of qPCR primers.

### RNA Sequencing

RNA was extracted as indicated above. RNA quality was measured using the 2100 Bioanalyzer (Agilent). Libraries were prepared using the KAPA Hyper Prep kit (Roche) and sequenced (sequencing read length, 75bp) in an Illumina HiSeq 4000 system. Briefly, reads were aligned using to relevant reference genome (mouse Ensembl GRCm38 - release 89 for 3LL and human Ensembl GRCh38 – release 38 for human cell lines). For data analysis, the R package DESeq2 was used and Gene Set Enrichment analysis was performed following gene sets available from MSigDB (Broad Institute).

### Whole exome sequencing and neoantigen prediction

DNA was extracted from cells using the QuickExtract DNA Extraction Solution (Lucigen) and sequencing was performed with 110x coverage using 100 base pair paired end read lengths. DNA library prep was performed using aSureSelectXT reagent kit (Agilent) and gDNA was sequenced using an Illumina HiSeq system.

Sequencing reads were aligned to the Mus musculus reference genome (mouse Ensembl GRCm38 - release 89). For mutation calling, DNA from wild-type C57Bl/6 mice was taken as a reference and analysed using the Mutect algorithm developed by the Broad Institute. Whole exome sequencing data of non-synonymous SNP-containing genes (in .vcf format) was combined with RNA sequencing data of expressed genes (TPM >0). Peptide sequences for obtained variants were converted using the SeqTailor tool from Rockefeller University (http://shiva.rockefeller.edu/SeqTailor/), by selecting the Mouse reference genome and a window size of 12aa on both sides of the variant. MHC binding prediction was performed using the IEDB 2.22 prediction method (http://tools.iedb.org/mhci/).

### Ex vivo immune cell culture and transwell assay

Femurs and tibias from C57Bl/6 mice were dissected and flushed using ice cold PBS using 21g needles. Flushed cells were centrifuged, filtered through a 45μM mesh and monocytes were magnetically isolated using the Monocyte Isolation Kit (BM, mouse) from Miltenyi as per manufacturer’s instructions.

Cell migration was quantified in duplicate using 24-well Transwell inserts (6.5mm) with polycarbonate filters (5μm pore size) (Corning Costar, Acton, MA). Monocytes (0.5×10^6^ in 100μl of RPMI) were added to the upper chamber of the insert. The lower chamber contained 600μl of RPMI 1640 medium or filtered conditioned medium from tumour cells. The plates were incubated at 37°C in 5% CO_2_ for 1.5h and cells that had migrated into the lower chamber were harvested and counted using flow cytometry.

### Cytokine assays

Medium from cells was harvested and used in the Human Cytokine Array Kit (R&D Systems), as per manufacturer’s instructions. For detection of CCL2, CXCL9 and CXCL10, Human CCL2/MCP-1 DuoSet ELISA, Mouse CCL2/JE/MCP-1 DuoSet ELISA, Mouse CXCL9/MIG DuoSet ELISA and Mouse CXCL10/IP-10 DuoSet ELISA kits (from R&D Systems) were used, following manufacturer’s instructions.

### Immunohistochemistry

Tumour-bearing lungs were fixed in 10% NBF for 24 h followed by 70% ethanol. Fixed tissue was embedded in paraffin wax. Tissue sections were stained with haematoxylin and eosin, using standard methods. For immunohistochemistry staining, tissue sections were boiled in sodium citrate buffer (pH 6.0) for 15 min and incubated with the following antibodies for 1h: anti-Foxp3 (D6O8R, CST), anti-CD8 (4SM15, Thermo Scientific). Primary antibodies were detected using biotinylated secondary antibodies and detected by HRP/DAB. Slides were imaged using a Leica Zeiss AxioScan.Z1 slide scanner

### Flow cytometry

Mice were culled using schedule 1 methods and lungs dissected (one spleen was also dissected to use as single stain control). Tumours were dissected from the lungs and cut into small pieces before incubating in digestion solution (1mg/ml collagenase type I and 50U/ml DNase in HBSS buffer) at 37°C for 30min. After homogenisation, samples were filtered through a 70mM cell strainer, erythrocytes were shocked using ACK lysing buffer (Life Technologies) and samples were re-filtered through 70mM cell strainers. After washes in PBS, samples were stained with fixable viability dye eFluor780 (BD HorizonTM) for 30min at 4°C. Samples were washed three times in FACS buffer (2mM EDTA, 0.5% BSA in PBS pH7.2) and stained using appropriate antibody mixes or single stain controls (spleen or OneComp eBeadsTM from ThermoFisher). After staining, samples were fixed in fix/lyse (Thermofisher) or FixPerm solution (Thermofisher) if intracellular staining was needed. Samples were then either stained with an intracellular antibody or washed and analysed using a FACSymphonyTM analyser (BD). Data was analysed using FlowJo software v10 (LLC).

For FACS analysis *in vitro*, cells were harvested with trypsin, filtered and washed in FACS buffer before appropriate antibody treatment. For intracellular cytokine staining, cells were treated with Brefeldin A (BD GolgiPlugTM) 6h before harvesting. Cells were permeabilised using the FixPerm (ThermoFisher) solution prior to staining. Samples were run in a LSRII or LSRFortessa (BD) and FlowJo software v10 (LLC) was used to analyse the data. For a list of antibodies used, see table 2.

**Table 2.** List of FACS antibodies.

| Target | Fluorophore | Clone | Source | Catalog number |
| --- | --- | --- | --- | --- |
| **H-2L<sup>d</sup>/H-2D<sup>b</sup>** | PE | 28-14-8 | Biolegend® | 114507 |
| **H2-Kb** | AF647 | AF6-88.5 | Biolegend® | 116512 |
| **CD45** | PerCP | 30-F11 | Biolegend® | 103130 |
| **CD3** | FITC | 17A2 | Biolegend® | 100204 |
| **gdTCR** | BV605 | GL3 | Biolegend® | 118129 |
| **CD4** | BUV737 | GK1.5 | BD Biosciences | 564298 |
| **CD8** | BUV395 | 53-6.7 | BD Biosciences | 563786 |
| **Foxp3** | eFluor660 | FJK-16s | eBioscience | 50-5773-82 |
| **CD44** | BV421 | IM7 | Biolegend® | 103039 |
| **CD62L** | BV711 | MEL-14 | Biolegend® | 104445 |
| **CD69** | BV605 | JES5-16E3 | Biolegend® | 104530 |
| **PD1** | BV785 | 29F.1A12 | Biolegend® | 135225 |
| **LAG3** | PE-Cy7 | eBioC9B7W | eBioscience | 25-2231-82 |
| **NKp46** | BV421 | 29A1.4 | Biolegend® | 137612 |
| **CD49b** | AF488 | DX5 | Biolegend® | 108913 |
| **CD19** | PE | 6D5 | Biolegend® | 115507 |
| **B220/CD45R** | BV605 | RA3-6B2 | Biolegend® | 103244 |
| **CD11c** | BUV395 | HL3 | BD Biosciences | 564080 |
| **CD11b** | BUV737 | M1/70 | BD Biosciences | 564443 |
| **Ly6G** | BV711 | 1A8 | Biolegend® | 127643 |
| **Ly6C** | BV785 | HK1.4 | Biolegend® | 128041 |
| **PD-L1** | PE | MIH5 | eBioscience | 12-5982-81 |
| **F4/80** | BV785 | EMR1 | Biolegend® | 123141 |
| **CD24** | BV605 | M1/69 | Biolegend® | 101827 |
| **CD103** | BV421 | M290 | BD Biosciences | 562771 |
| **CD64** | PE-Cy7 | X54-5/7.1 | Biolegend® | 139314 |
| **CD206** | BV711 | C068C2 | Biolegend® | 141727 |
| **TIM3** | PE | RMT3-23 | Biolegend® | 119703 |
| **CD86** | BV785 | GL-1 | Biolegend® | 105043 |
| **MHCII** | FITC | M5/114.15.2 | Biolegend® | 107605 |
| **CXCL9** | PE | MIG-2F5.5 | Biolegend® | 515603 |

### Imaging Mass Cytometry

Tissue processing and antibody staining was performed as described in detail in (21). In short, 5µm cryosections of fresh frozen lungs were fixed (Image-iT™ Fixative Solution, ThermoFisher) and stained with the antibody panel listed in Table 3 and Cell-ID Intercalator-Ir (Fluidigm). Scanning of the (dried) slides was done with the Hyperion Imaging Mass Cytometer (Fluidigm). Images available upon request.

**Table 3.** List of IMC antibodies.

| **Metal** | **Target** | **Clone** | **Source** | **Catalog no.** |
| --- | --- | --- | --- | --- |
| **89Y** | CD45 | 30-F11 | Fluidigm | 3089005B |
| **141Pr** | aSMA | 1A4 | Fluidigm | 3141017D |
| **142Nd** | MHCcII | M5/114.15.2 | BioLegend | 107637* |
| **144Nd** | MHCcI | 28-14-8 | Fluidigm | 3144016C |
| **146Nd** | F480 | (Cl:A3-1) | BioRAD | MCA497GA* |
| **147Sm** | CD68 | FA-11 | BioLegend | 137002* |
| **150Nd** | CD44 | IM7 | Fluidigm | 3150018B |
| **152Sm** | CD3e | 145-2C11 | Fluidigm | 3152004B |
| **153Eu** | PDL1 | 10F.9G2 | Fluidigm | 3153016B |
| **158Gd** | Foxp3 | FJK-16s | Fluidigm | 3158003A |
| **161Dy** | CD103 | AF1990G | R&D systems | AF1990* |
| **165Ho** | TIGIT | 4D4/mTIGIT | BioLegend | 156102* |
| **166Er** | PD1 | 29F.1A12 | BioLegend | 135202* |
| **167Er** | NKp46 | 29A1.4 | Fluidigm | 3167008B |
| **168Er** | CD8a | 53-6.7 | Fluidigm | 3168003B |
| **169Tm** | CD4 | RM4-5 | BioLegend | 100561* |
| **170Er** | CXCL9 | MIG-2F5.5 | BioLegend | 515602* |
| **171Yb** | Granzyme B | GB11 | Fluidigm | 3171002C |
| **172Yb** | cleaved caspase<br>3 | 5A1E | Fluidigm | 3172027D |
| **173Yb** | Ki67 | 16A8 | BioLegend | 652402* |
| **174Yb** | LAG3 | C9B7W | Fluidigm | 3174019C |
| **176Yb** | B220 | RA3-6B2 | Fluidigm | 3176002B |
| **209Bi** | CD11c | N418 | Fluidigm | 3209005B |

\*Antibodies conjugated in house using MaxPar antibody labelling kits (Fluidigm).

Image processing was performed with the previously described 1px-expansion single cell segmentation pipeline using imcyto (nf-core/imcyto). The resulting single cell data was clustered with Phenograph, and subsequently annotated to the different cell types (Supplementary Table 1).

### Statistical analysis

For most experiments, data were compared using unpaired or paired two-tailed Student’s t-tests, or ANOVA if more than two experimental groups were examined. In mouse tumour analysis, the Mann-Whitney u-test was used for volume comparison. To compare read counts of individual genes in mRNA-Seq datasets of two groups, Wald test was used with a Benjamini and Hochberg correction with an FDR Q value of 5% to obtain adjusted p values (Statistical analysis was performed by Crick Bioinformatics Facility). To compare two survival curves, the Mantel-Cox log-rank test was used. Statistical analyses were performed in Prism 7 (GraphPad Software) or in RStudio. Significance is presented as *P < 0.05, **P < 0.01, ***P < 0.001 and ****P < 0.0001.

## Acknowledgments

We thank the science technology platforms at the Francis Crick Institute including the Biological Research Facility, Advanced Sequencing Facility, Computational Biology, Genomics Equipment Park, Experimental Histopathology, FACS and Cell Services. We thank Philip Hobson and Dina Levi for assistance with imaging mass cytometry.

## Author Contributions

E.M, F.vM., JB, M.M-A., and J.D. designed the study, interpreted the results and wrote the manuscript. E.M, F.vM, M.M-A., S.R., J.B., R.A., P.A., P.R.C., K.V. performed the biochemical experiments, C.M. assisted with in vivo studies, M.L.S. and P.E. performed bioinformatics analyses. All authors contributed to manuscript revision and review.

## Data and materials availability

All data needed to evaluate the conclusions in the paper are present in the paper and/or the Supplementary Materials. RNA sequencing data has been deposited at Gene Expression Omnibus (accession codes GSE183549 and GSE199582). Imaging mass cytometry raw images (figure 4E) are available at the Figshare repository https://doi.org/10.25418/crick.19590259.

## SUPPLEMENTAL MATERIAL

**Fig. S1.**
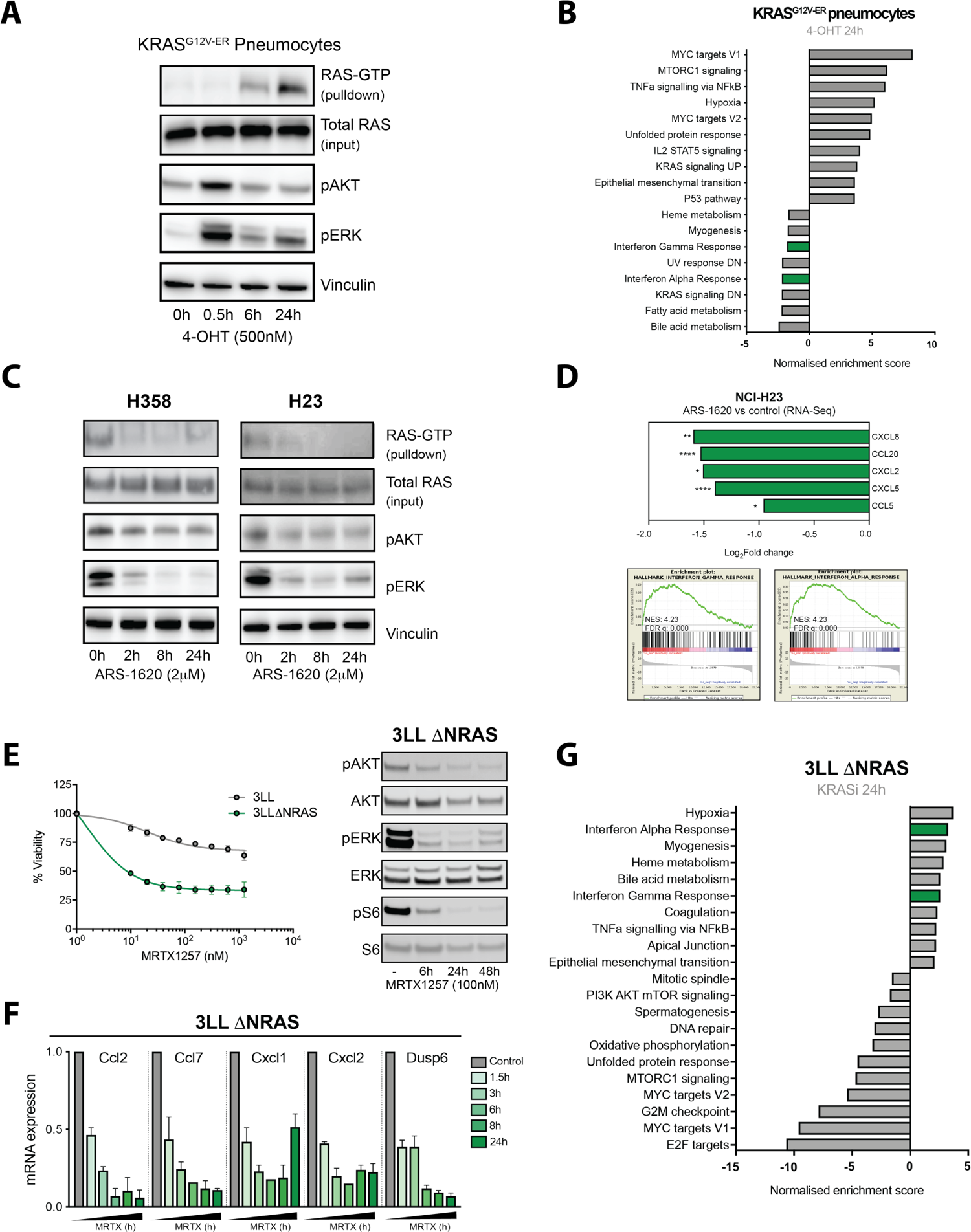
KRAS-dependent regulation of secreted factors in human and murine cell lines. (A) Time course of KRAS^G12V-ER^ pneumocytes treated with 500nM 4-OHT showing increased active KRAS (KRAS-GTP) and downstream pathway activation. (B) Summary of most up- and down-regulated pathways (MSigDB Hallmarks, FDR q<0.05) in 4-OHT treated KRAS^G12V-ER^ pneumocytes. (C) Time course of human KRAS^G12C^ lung cancer cell lines treated with 2μM ARS-1620 showing decreased active KRAS (KRAS-GTP) and downstream pathway activation. (D) Top: Log_2_Fold change of selected cytokine genes from RNA-Seq data in ARS-1620 (2μM, 24h, p adjusted value) treated NCI-H23 cells versus DMSO control. Bottom: MSigDB Hallmarks GSEA plots of IFN*α* and IFN*γ* pathway genes in ARS-1620 treated versus control samples. (E) Left: viability assay comparing parental 3LL and CRISPR-edited 3LL *Δ*NRAS cells treated with increasing concentrations of MRTX1257 for 72h (n=2 independent experiments, mean±SEM). Right: Time course of 3LL *Δ*NRAS cells treated with 100nM MRTX1257 showing downstream pathway inhibition. (F) Time course analysis of MRTX1257-treated 3LL *Δ*NRAS cells showing mRNA expression of cytokines and Dusp6 as a control for KRAS inhibition (2^-ΔΔCT^, normalised to control sample for all genes, n=2, mean+SEM). (G) Summary of most up- and down-regulated pathways (MSigDB Hallmarks, FDR q<0.05) in ARS-1620-treated 3LL *Δ*NRAS cells.

**Fig. S2.**
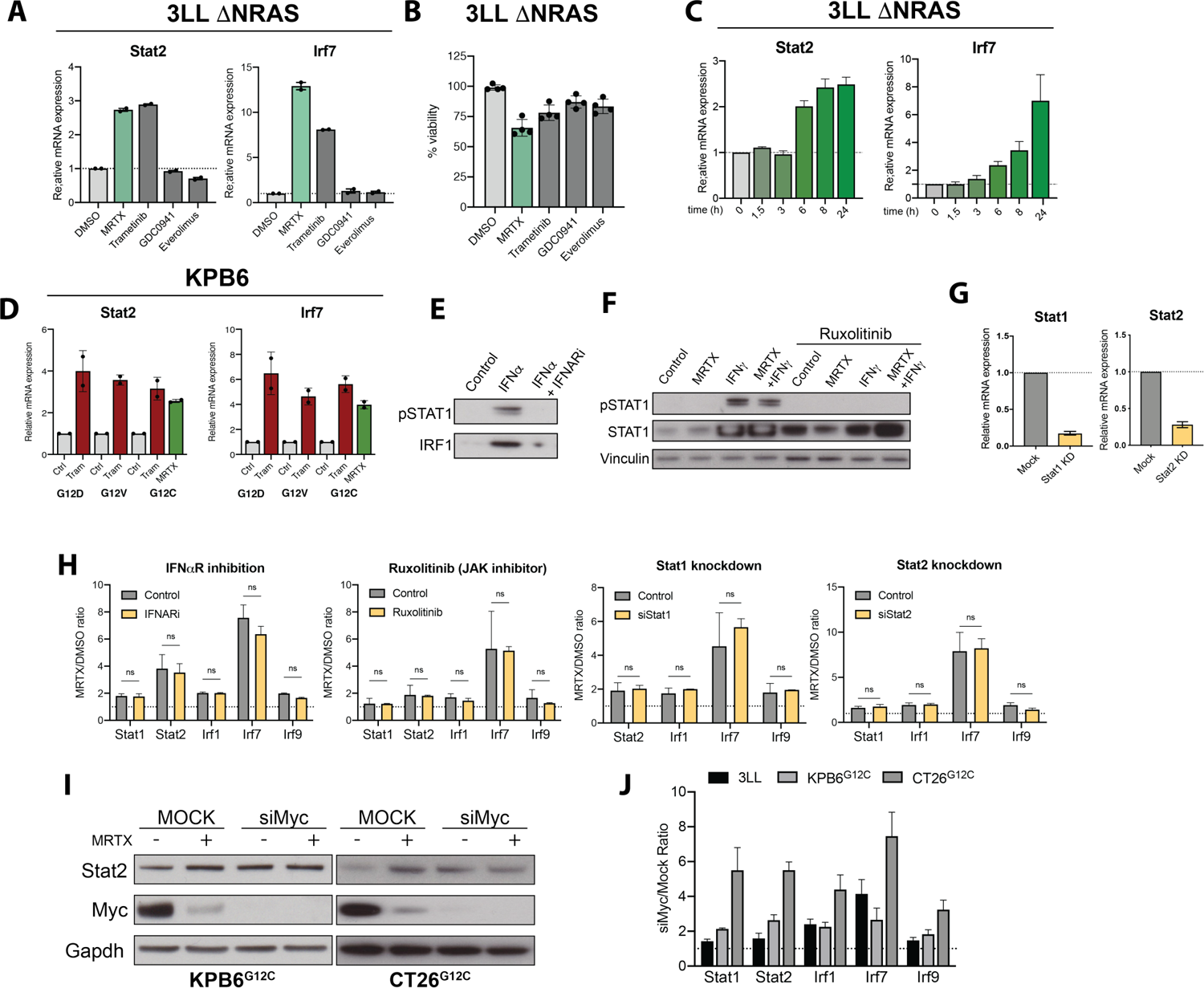
Mechanism of KRAS-dependent regulation of IFN genes. (A) Expression of IFN-induced genes *Stat2* and *Irf7* in 3LL *Δ*NRAS cells treated with DMSO control, MRTX1257 (100nM), MEK inhibitor trametinib (10nM), PI3K inhibitor GDC0941 (500nM) or mTOR inhibitor everolimus (100nM) for 24h (2^-ΔΔCT^, normalised to control sample, n=2, mean±SEM). (B) Viability of 3LL *Δ*NRAS cells treated with drugs as in (A) for 24h (n=4). (C) 100nM MRTX1257 treatment time course of 3LL *Δ*NRAS cells (2^-ΔΔCT^, normalised to control sample, n=3, mean+SEM). (D) mRNA expression of Stat2 and Irf7 after trametinib (10nM, 24h) treatment of isogenic KPB6 cell lines with differing KRAS G12 mutations and comparison with MRTX1257 (100nM) treatment in KPB6^G12C^ cell line. (2^-ΔΔCT^, normalised to control sample for each cell line, n=2, mean±SEM) (E) Western blot showing loss of IFN sensitivity after IFN*α*R (20mg/ml, 24h) blocking antibody treatment (100ng/ml IFN*α*, 24h) of 3LL *Δ*NRAS cells. (F) Western blot showing loss of IFN sensitivity by 1μM ruxolitinib (100ng/ml IFN*γ*, 24h). (G) Knockdown efficiency of siStat1 (48h, left) and siStat2 (48h, right) measured by qPCR (2^-ΔΔCT^, normalised to control sample, n=3). (H) Comparison of the ratio of IFN-induced gene expression in MRTX-versus DMSO-treated 3LL *Δ*NRAS in control cells and cells treated with an anti-IFNaR antibody treatment (20mg/ml), 1μM JAK1/2 inhibitor Ruxolitinib treatment, Stat1 or Stat2 knockdown (24h, 2^-ΔΔCT^, n=3, paired t test). (I) Western blot of KPB6^G12C^ and CT26^G12C^ cells showing *Myc* knockdown and *Stat2* increase after treatment with MRTX, Myc siRNA, or both. (J) Ratio of Myc siRNA versus mock control treatment expression of IFN-induced genes in 3LL *Δ*NRAS (n=3), KPB6^G12C^ (n=4) and CT26^G12C^ (n=3) cells.

**Fig. S3.**
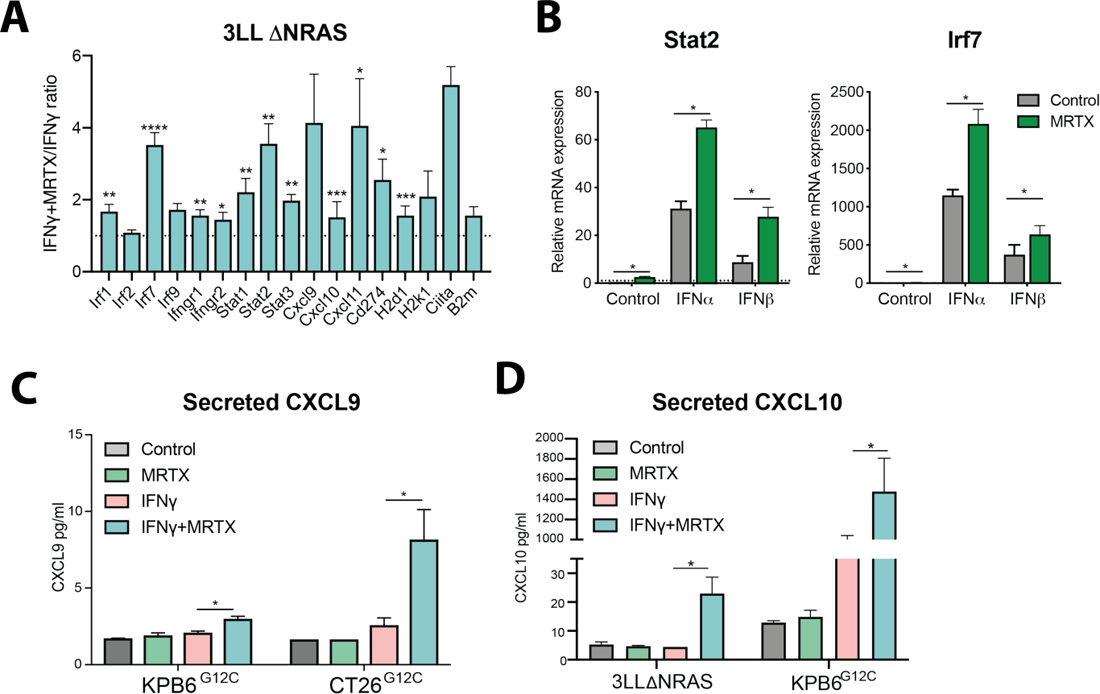
KRAS-driven augmentation of type I and II IFN treatment response. (A) Summary of all IFN-stimulated genes (ISG) examined, showing the ratio of IFN*γ* plus MRTX1257 (100nM) versus IFN*γ* (100ng/ml) alone (24h treatment, 2^-ΔΔCT^, normalised to IFN*γ*-treated sample for all genes, n at least 3, mean+SEM). (B) mRNA expression of IFN-induced genes *Stat2* and *Irf7* after treatment of 3LL *Δ*NRAS cells with recombinant IFN*α*/*β*(100ng/ml) and/or MRTX for 24h (2^-ΔΔCT^, normalised to control, n=3, paired t test, mean+SEM). (C) Concentration of CXCL9 secreted to the cell culture supernatant by CT26^G12C^ and KPB6^G12C^ cells after treatment with MRTX, IFN*γ* or both (normalised to control, n=2 for KPB6^G12C^, n=3 for CT26^G12C^, mean+SEM, paired t test). (D) Concentration of CXCL10 secreted to the cell culture supernatant by 3LL *Δ*NRAS and KPB6^G12C^ cells after treatment with MRTX, IFN*γ* or both (normalised to control, n=3, mean+SEM, paired t test).

**Fig. S4.**
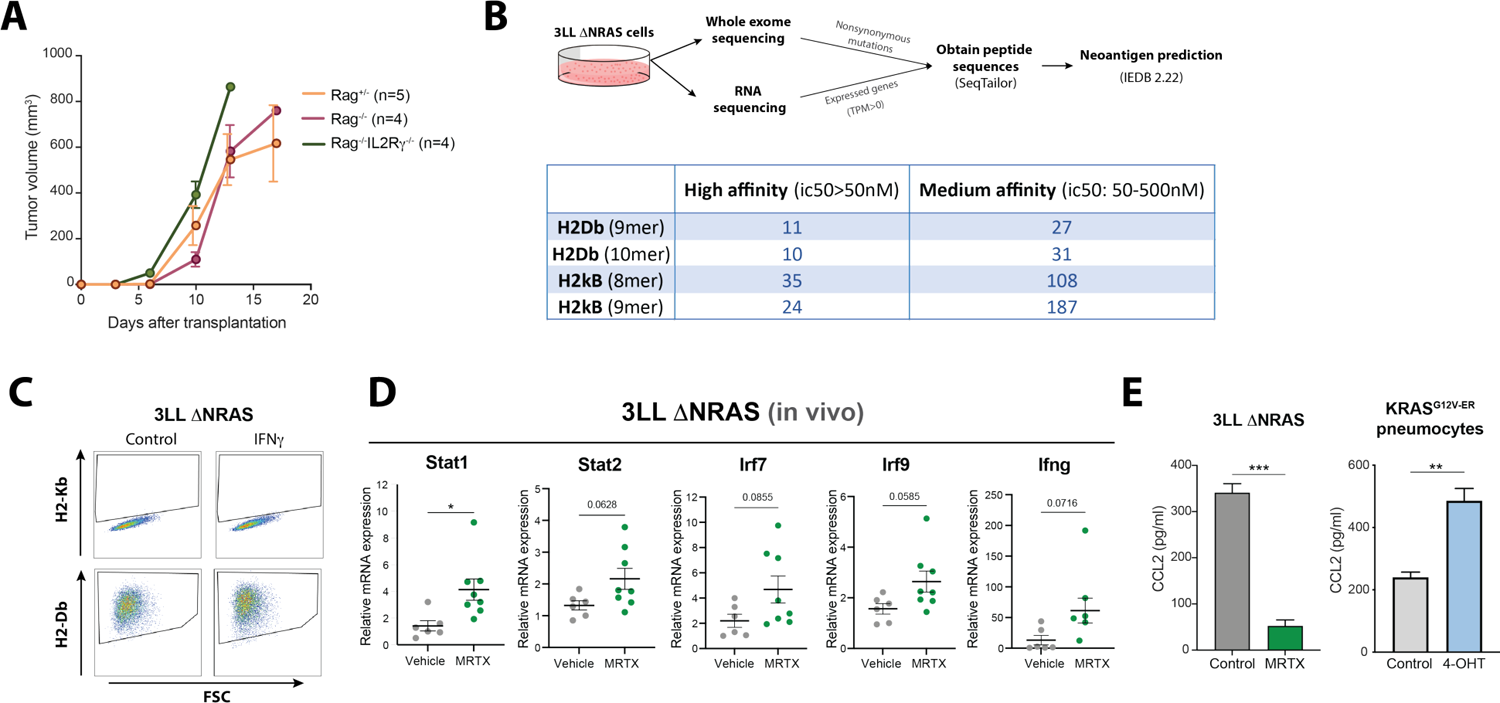
In vivo characterisation of 3LL ΔNRAS tumours and the effects of KRAS^G12C^ inhibition on myeloid cells. (A) Growth comparison of subcutaneously implanted 3LL *Δ*NRAS tumours in Rag1^+/-^, Rag1^-/-^ and Rag1^-/-^IL2R*γ*^-/-^ mice. (B) Above: summary of in silico analysis merging whole exome sequencing and RNA-Seq data to obtain predicted neoantigens. Below: number of predicted high and medium affinity neoantigens obtained for each C57Bl/6 MHC allele for different sized peptides. (C) Flow cytometric analysis of 3LL *Δ*NRAS cells in vitro showing lack of basal and IFN*γ*-induced (100ng/ml, 24h) expression of surface H2-Kb, and intact H2-Db expression. (D) qPCR analysis of IFN-induced genes in 3LL *Δ*NRAS lung tumours (2^-ΔΔCT^, vehicle n=6, MRTX n=8, unpaired t test, mean±SEM). (E) Concentration of secreted CCL2 as measured by ELISA in medium from either control or 4-OHT-treated KRAS^G12V-ER^ pneumocytes (n=3, Mean+SEM, unpaired t test).

**Fig. S5.**
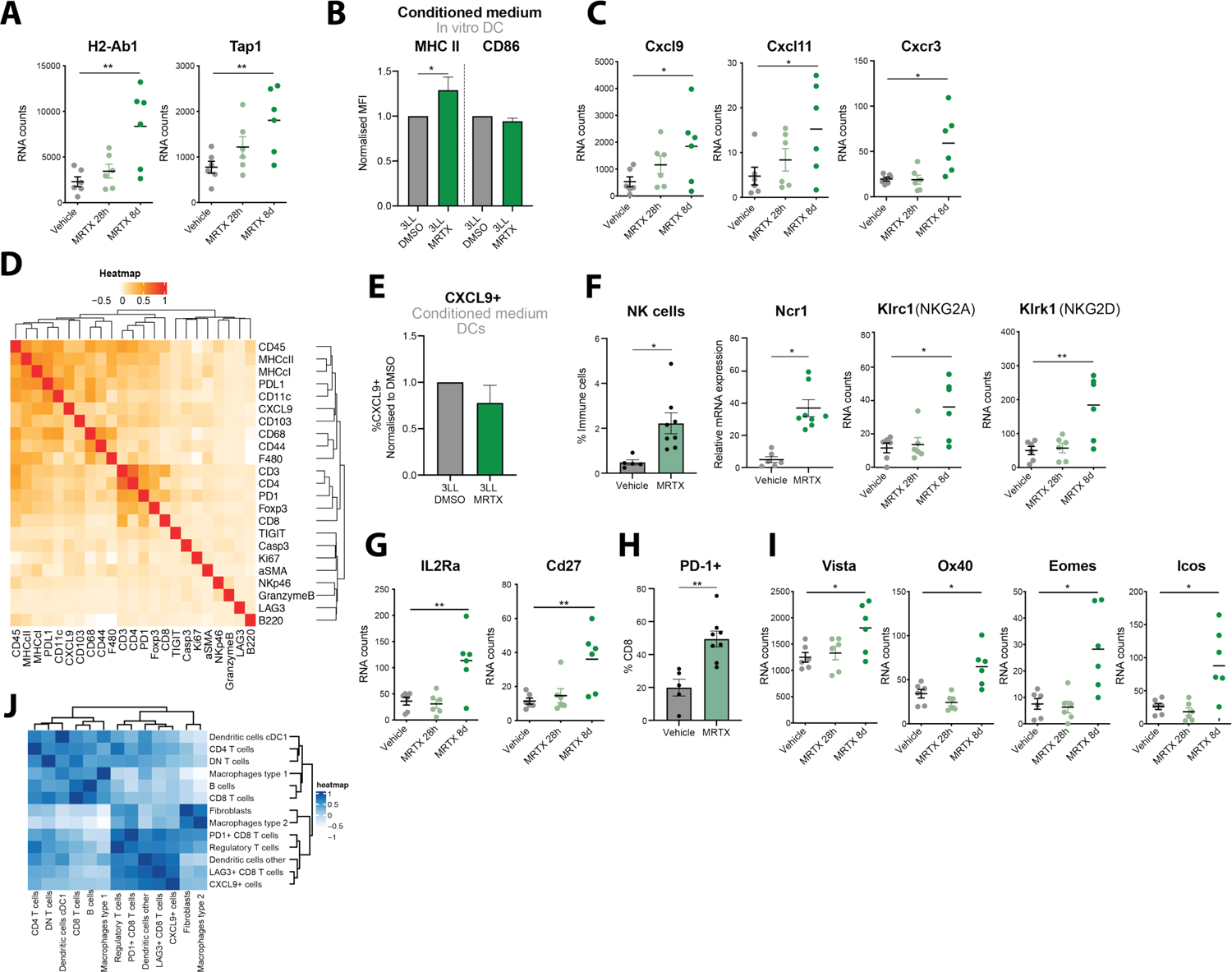
KRAS^G12C^ effects on antigen presentation, T cell infiltration and activation in vivo. (A) mRNA counts showing increased expression of antigen presentation genes from RNA-Seq analysis of 3LL *Δ*NRAS lung tumours treated with vehicle or 50mg/kg MRTX1257 for 28h or 8d (FDR p adjusted value, n=6 tumours per group). (B) Normalised mean fluorescence intensity of MHC II and CD86 on DCs cultured overnight under filtered conditioned medium from either DMSO-treated or MRTX-treated (48h) 3LL *Δ*NRAS cells (n=3, mean+SEM, 2-way ANOVA). (C) mRNA counts for T cell chemoattractant and receptor-encoding genes, analysed as in (A). (D) Pearson correlation matrix of markers expressed at single cell level as measured by IMC. (E) Normalised percentage of CXCL9+ DCs after overnight incubation as in (B), analysed as in (B). (F) NK cell data summary after one week of MRTX1257 treatment in vivo. Left: increased NK cell infiltration in tumours as measured by flow cytometry (pre-gated as CD45+ CD19-NKp46+ CD49b+, n=5 for vehicle, n=8 for MRTX-treated, unpaired t test). Middle: qPCR analysis for NK cell marker Ncr1 (6 samples per group, unpaired t test). Right: mRNA count data for NK cell markers *Klrc1* and *Klrk1*, analysed as in (A). (G) mRNA counts for T cell activation genes *IL2Rα* and *Cd27*, analysed as in (A). (H) Percentage of PD-1+ CD8+ T cells measured by flow cytometry (vehicle n=5, MRTX n=8, unpaired t test, mean±SEM). (I) mRNA counts showing increased expression of T cell exhaustion genes from RNA-Seq of lung 3LL *Δ*NRAS tumours treated with vehicle or MRTX, analysed as in (A). (J) Pearson correlation matrix based on cell proportions present within the tumour and interface domain of MRTX-treated tumours measured by IMC.

**Fig. S6.**
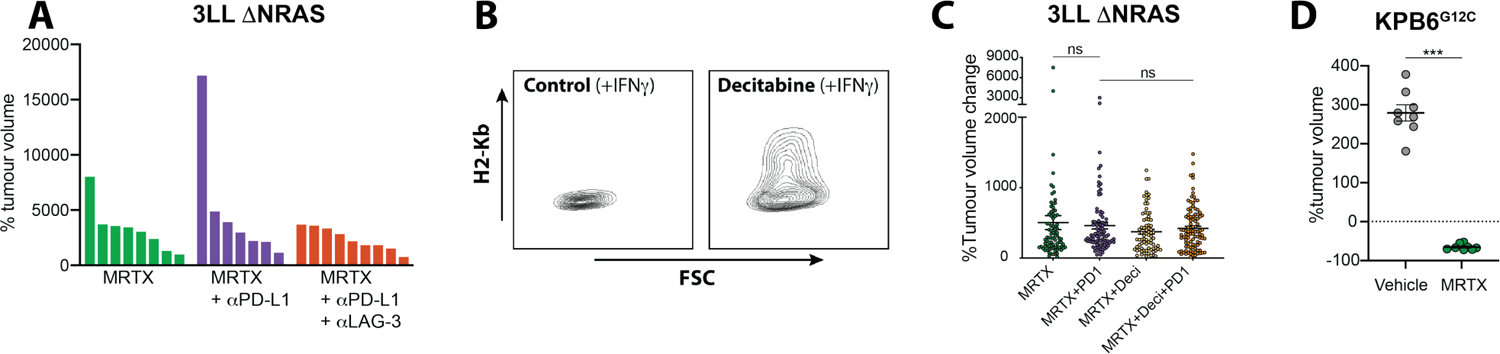
Lack of combinatorial effects of KRAS^G12C^ inhibition and ICB in immune refractory tumours. (A) Waterfall plot showing tumour volume change of 3LL *Δ*NRAS lung tumours treated with MRTX only (n=8 mice), 50mg/kg MRTX1257 and 10mg/kg anti-PD-L1 antibody (n=7 mice) or 50mg/kg MRTX1257, 10mg/kg anti-PD-L1 and 10mg/kg anti-LAG3 antibody combination (n=9) displaying no combinatorial effect. (B) H2-kB expression measured by flow cytometry of IFNg (100ng/ml) treated 3LL *Δ*NRAS cells after decitabine (5’Aza-2’-deoxycytidine, 250nM, 24h) treatment. (C) Tumour volume change of 3LL *Δ*NRAS lung tumours treated with MRTX1257 (50mg/kg, n=9 mice), MRTX+anti-PD-1 (10mg/kg, n=10 mice), MRTX+Deci (0.3mg/kg, n=9 mice) or the triple combination (7 day treatment, 2-way ANOVA). (D) Tumour volume change of KPB6^G12C^ tumours after one week treatment with 50mg/kg MRTX1257 showing marked regression (n=8 mice per group, Mann-Whitney analysis).

**Fig. S7.**
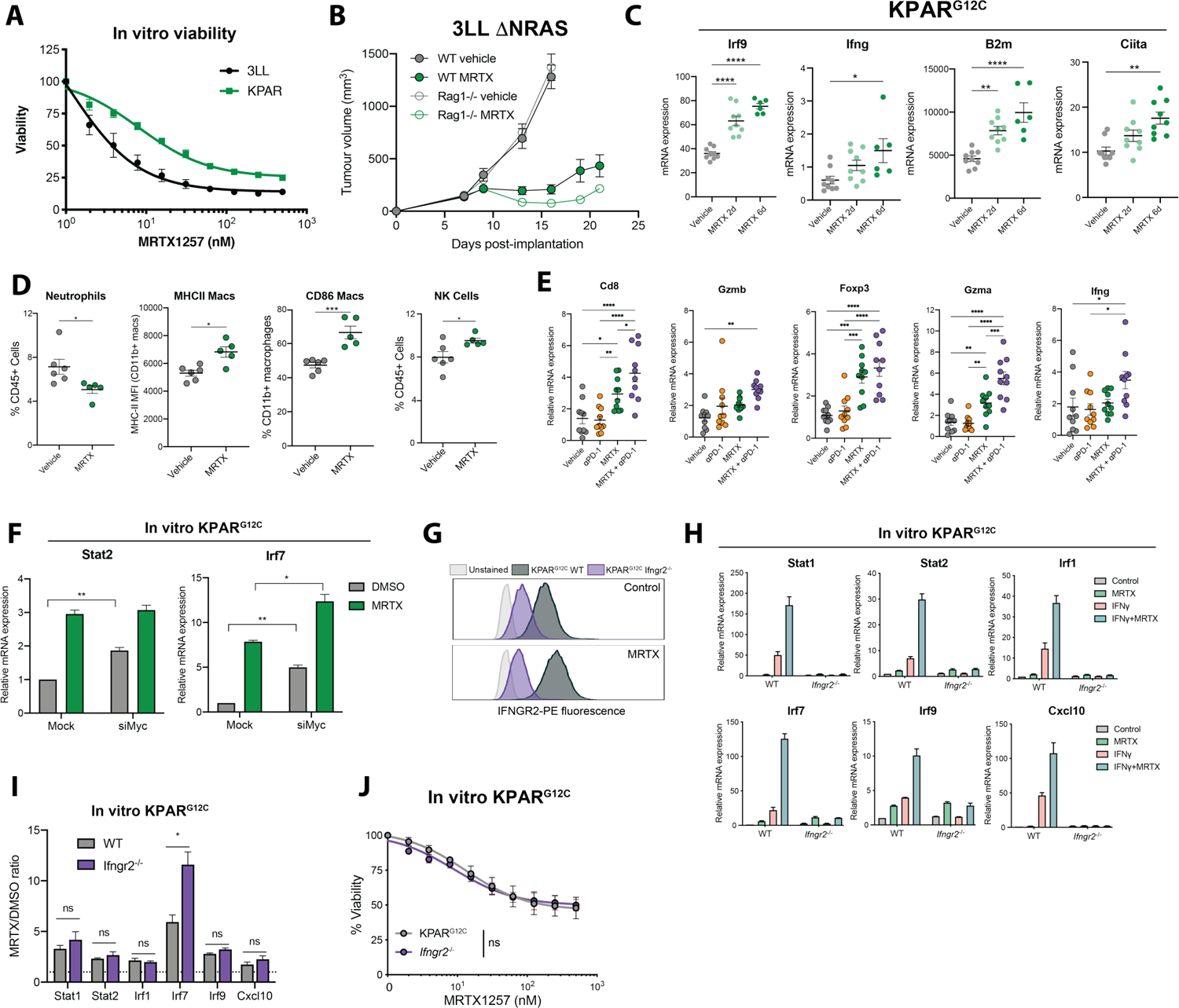
Effects of KRAS^G12C^ inhibition in immunogenic tumours and role of IFN signalling. (A) Viability data for 3LL *Δ*NRAS and KPAR^G12C^ cells treated in vitro with increasing concentrations of MRTX1257 for 72h. (B) Subcutaneous 3LL *Δ*NRAS tumour growth in C57Bl/6 WT mice treated with vehicle (n=6 mice) or MRTX1257 50mg/kg (n=7 mice) and *Rag1*^-/-^ mice treated with vehicle (n=9 mice) or MRTX1257 (n=10 mice). (C) qPCR analysis of IFN-induced and antigen presentation genes in KPAR^G12C^ tumours treated with vehicle (n=9 tumours) or 50mg/kg MRTX849 for two (n=9 tumours) or 6 days (n=6 tumours, one way ANOVA). (D) Flow cytometry data from MRTX849 treated (4 days) KPAR^G12C^ lung tumours. Neutrophils are gated as live CD45+ CD11b+ Ly6C+ Ly6G+, macrophages as live CD45+ CD11b+ CD24-CD64+ and NK cells as Live CD45+ CD19-NKp46+ CD49b+ (control n=6 mice, MRTX n=5 mice, unpaired t test). (E) qPCR analysis of immune markers on KPAR^G12C^ lung tumours treated for 5 days with 50mg/kg MRTX849 and/or 10mg/kg anti-PD-1, n=10 mice per group (each dot represents a tumour, ANOVA multiple comparisons test). (F) qPCR analysis of IFN-induced genes KPAR^G12C^ cells treated with MRTX1257, Myc siRNA or both (2^-ΔΔCT^, normalised to control sample for all genes, n=3, paired t tests siMyc versus Mock, mean+SEM). (G) Flow cytometry analysis of surface IFNGR2 expression on KPAR^G12C^ or *Ifngr2*^-/-^ cells in vitro, treated with DMSO or 100nM MRTX1257 for 24h. (H) In vitro qPCR analysis of KPAR^G12C^ WT and *Ifngr2*^-/-^ cells treated with 100nM MRTX1257, 100ng/ml IFN*γ* or both for 24h (2^-ΔΔCT^, paired t test, n=3, mean+SEM). (I) Same data as in (H), showing the MRTX/DMSO ratio of expression of IFN-related genes in KPAR^G12C^ WT vs *Ifngr2*^-/-^ cells (unpaired t test). (J) In vitro viability of WT and *Ifngr2*^-/-^ KPAR^G12C^ cells treated with a range of doses of MRTX1257 for 72h (n=3, mean±SEM).

---

**Supplementary table 1. IMC single cell data**

Single cell data, obtained by segmentation and clustering, of the imaging mass cytometry image dataset deposited on Figshare (https://doi.org/10.25418/crick.19590259).

Each row provides the data for a single cell. Column description:

- “ROI_name” gives the name of the image file as deposited on Figshare.
- “MouseID”, code for the mouse from which the tumour was originally taken.
- “treatment” lists whether a tumour was treated with Vehicle or MRTX.
- “MI_” denotes mean intensity per cell for the markers listed.
- “cluster” the number of cluster assigned by Phenograph and refined supervised gating.
- “clustername” the cell type that was manually assigned based on expression profile.
- “Location_Center_X/Y” represent the coordinates of the centre of each cell in the image.
- “dist_” lists the distances for every cell in the image to the nearest of the cell type as called in the name of the column.

